# Differentiation SELEX approach identifies RNA aptamers with different specificities for HIV-1 capsid assembly forms

**DOI:** 10.1101/2023.12.11.571135

**Authors:** Paige R. Gruenke, Miles D. Mayer, Rachna Aneja, Zhenwei Song, Donald H. Burke, Xiao Heng, Margaret J. Lange

**Author notes:** Margaret Lange, Department of Molecular Microbiology & Immunology, University of Missouri School of Medicine, Columbia, Missouri, 65212, United States.

## Abstract

The HIV-1 capsid protein (CA) assumes distinct assembly forms during replication, each presenting unique, solvent-accessible surfaces that facilitate multifaceted functions and host factor interactions. However, contributions of individual CA assemblies remain unclear, as the evaluation of CA in cells presents several technical challenges. To address this need, we sought to identify CA assembly form-specific aptamers. Aptamer subsets with different specificities emerged from within a highly converged, pre-enriched aptamer library previously selected to bind the CA hexamer lattice. Subsets were either highly specific for CA lattice or bound both CA lattice and CA hexamer. We further evaluated four representatives to reveal aptamer structural features required for binding, highlighting interesting features and challenges in aptamer structure determination. Importantly, our aptamers bind biologically relevant forms of CA and we demonstrate aptamer-mediated affinity purification of CA from cell lysates without virus or host modification. Thus, we have identified CA assembly form-specific aptamers that represent exciting new tools for the study of CA.

## INTRODUCTION

HIV-1 capsid protein (CA) assembly forms are involved in multiple viral replication steps including assembly, maturation, reverse transcription, cytoplasmic trafficking, nuclear import, capsid uncoating, and integration (*1–7*). CA is present in the Gag and GagPol polyproteins during virus assembly and as part of the immature Gag lattice (*8–11*). Upon budding and activation of the viral protease, Gag and GagPol are cleaved into their component parts and newly liberated CA monomers assemble through various intermediates into the mature capsid core composed of hexamers and exactly 12 pentamers (*12*). The mature core is released into the cytoplasm following viral fusion, where it utilizes cellular trafficking machinery to reach the nucleus (*2, 3, 13–16*). During trafficking, the capsid core supports initiation of reverse transcription, nuclear entry, completion of reverse transcription, integration, and evasion of the host immune response (*7, 17–21*). Due to the many roles of CA in replication, it is unsurprising that CA mutations that disrupt structure and/or stability have a significant impact on viral infectivity (*22–27*). Indeed, maintenance of structure and appropriate levels of stabilization within CA assemblies at each stage of replication are critical for allowing required host factor interactions and supporting productive infection.

Despite significant advances in our understanding of CA structure and function, there are many unresolved questions related to the mechanistic roles of CA in viral replication and the contributions of individual CA assembly states to both early and late stages of the replication, including those related to CA-host factor interactions. A significant barrier to the resolution of these questions is the genetic fragility of CA. Traditional affinity purification methods for host factor identification require modification of the protein of interest, but CA is not easily amenable to modification (*24, 28, 29*) or mutation due to significant impacts on infectivity and alteration of numerous replication stages (*22, 30–32*). Furthermore, as HIV often utilizes host factors important for cellular function (*29, 33*), eliminating or reducing expression of those factors can significantly impact cellular biology or result in cell death, precluding a complete mechanistic understanding of HIV-host factor interactions. Despite the importance of CA assembly forms, there are no tools available that discriminate among CA assemblies to enable identification and study of CA assembly form-specific interactions in cells. Better understanding the role and interactions of CA assemblies in viral replication may enable the identification of novel sites on CA amenable to therapeutic targeting.

Nucleic acid aptamers fold into unique structures capable of discriminating among proteins, including those that differ by only a single amino acid mutation (*34–36*) or different conformations of the same protein (*37–42*). Aptamers are selected through an iterative process known as Systematic Evolution of Ligands by EXponential enrichment (SELEX) (*43, 44*) and can be easily modified for diverse downstream applications. Further, RNA aptamers can be expressed in cells. Thus, it is feasible that aptamers selected to differentiate among CA assembly forms could be developed as biochemical tools to improve our understanding of the roles and interactions of CA assembly forms during viral replication. We recently described an RNA aptamer selection against the assembled HIV-1 CA hexamer lattice (*45*). While the selected libraries demonstrated dose-dependent binding to the CA hexamer lattice but not the soluble CA hexamer or CA monomer, we reasoned that other low-abundant sequences capable of binding other CA assembly forms may be present within the libraries (Figure 1A). Here, we describe a readily generalizable differentiation selection strategy that successfully identified subsets of aptamers from a highly converged, pre-enriched CA hexamer lattice-binding aptamer library that were either CA hexamer lattice-specific binders or capable of binding both CA hexamer lattice and soluble CA hexamer. Using a combination of bioinformatic and biochemical approaches, we then defined key sequence and structural features required for CA binding for four aptamers; two that are specific for the CA hexamer lattice and two that bind both CA hexamer lattice and soluble CA hexamer. Finally, we performed a preliminary proof-of-concept experiment demonstrating that these aptamers can be adapted for affinity purification of CA from cell lysates.

**Figure 1:**
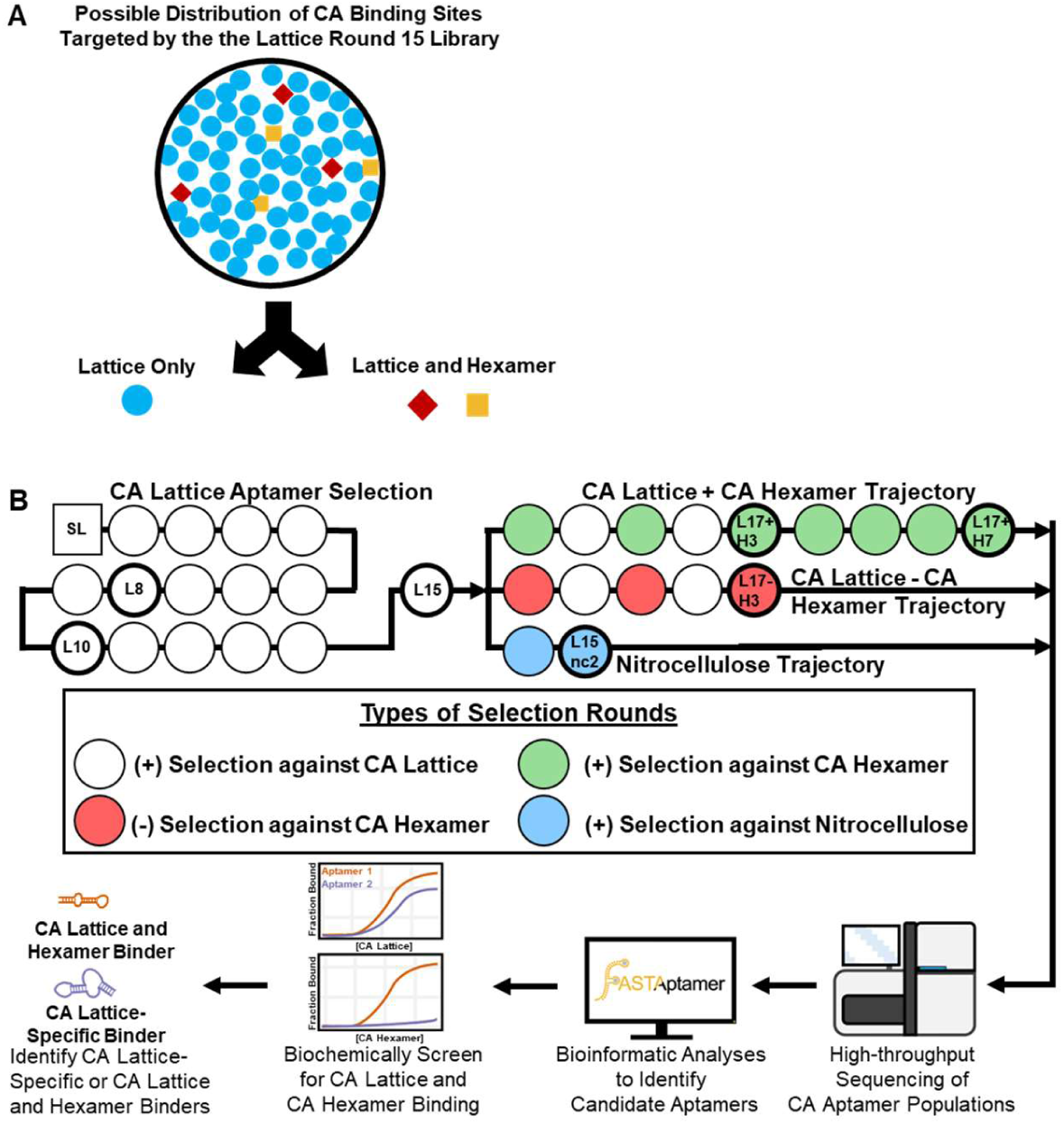
Capsid Differentiation Selection. (A) Schematic depicting the possible distribution of binding sites on the CA targeted by the lattice round 15 aptamer library (L15). Aptamers selected against the CA lattice could bind on sites present on each CA monomer (triangles), assembled CA hexamer (squares), or at hexamer-hexamer interfaces (circles). (B) Starting from L15, three selection trajectories were performed: the CA lattice and hexamer trajectory, the hexamer depletion trajectory, and the nitrocellulose trajectory. The seven libraries (labeled; bold, thick outlines) were submitted for high-throughput sequencing and candidate aptamers were identified for further characterization based on their enrichment and/or depletion profiles. Candidate aptamers were then screened for binding to CA lattice and hexamer to confirm CA lattice-specific or CA lattice and hexamer binders.

## MATERIAL AND METHODS

### Reagents

Unless otherwise noted, all chemicals were purchased from Sigma-Aldrich (St. Louis, MO). DNA templates for aptamers and all primers were ordered from Integrated DNA Technologies (Coralville, IA). The sequences of the aptamer RNAs and DNA primers used in this study are listed in Table S4.

Plasmid constructs for expression and purification of soluble HIV-1 CA hexamer (pET11a-CA (A14C, E45C, W184A and M185A)), CA hexamer lattice (pET11a-CA (A14C and E45C)), and CA monomer (pET11a-CA) were kindly provided by Dr. Owen Pornillos and Dr. Barbie Ganser-Pornillos (University of Utah).

### Expression and purification of HIV-1 CA proteins and crosslinking of CA hexamers and CA lattice tubes

HIV-1 CA protein assemblies were expressed and purified as previously described (*46, 74*). Briefly, mutant HIV-1 CA proteins were expressed using IPTG induction in *Escherichia coli* BL21(DE3) cells for 6–12 hr at 25°C. The monomeric p24 CA protein was precipitated using 25% ammonium sulfate and the pellet was resuspended in 20 mM Tris pH 8.0, 40 mM NaCl, and 60 mM βME, followed by purification using a Hi-Trap Q-Sepharose column. Purified protein fractions were concentrated and reconstituted in 25 mM Tris-HCl (pH 8.1) and 40 mM NaCl, flash frozen and stored at -80°C.

Soluble CA hexamers (pET11a-CA (A14C/E45C/W184A/M185A)) and CA lattice tubes (pET11a-CA (A14C/E45C)) were assembled as previously described (*46, 75*). Briefly, assembly was performed *in vitro* using sequential overnight dialysis. Pooled and concentrated fractions were dialyzed into assembly buffer (50 mM Tris, pH 8.1, 1.0 M NaCl, 200 mM βME) followed by dialysis into storage buffer (20 mM Tris, pH 8.1, 40 mM NaCl). The integrity of the assembled CA lattice tubes was verified using transmission electron microscopy (TEM) with assistance from the University of Missouri Electron Microscopy Core. The purity and integrity of monomeric p24, assembled soluble CA hexamers and CA lattice were confirmed using non-reducing SDS-PAGE. Importantly, for the purposes of this study, all CA protein concentrations are reported in terms of the total CA monomer.

### CA Differentiation Selection

The starting library for this work was the HIV-1 assembled CA lattice round 15 library described previously (*45*). To initiate the differentiation selection, double-stranded DNAs from this library were first PCR-amplified using Taq DNA polymerase for 25 cycles and then transcribed using the Y639F mutant T7 RNA polymerase purified in house (*76*), *in vitro* transcription buffer (50 mM Tris-HCl pH 7.5, 15 mM MgCl_2_, 5 mM DTT, 2 mM spermidine), and 2 mM of each NTP (ATP, CTP, GTP, and UTP). Reactions were incubated at 37°C for a minimum of 4 hr and halted by adding an equal volume of denaturing gel loading buffer (90% formamide and 50 mM EDTA with trace amounts of xylene cyanol and bromophenol blue). RNAs were purified using denaturing PAGE (6% TBE-PAGE, 8 M urea). Bands corresponding to the expected product sizes were visualized by UV shadow, excised, and eluted overnight in 300 mM sodium acetate (pH 5.4) at room temperature. Eluates were ethanol precipitated, resuspended in nuclease-free water and stored at -20°C. RNA concentrations were determined using a NanoDrop One spectrophotometer (ThermoFisher Scientific, Waltham, MA).

For each selection trajectory, the order and types of selection rounds performed are shown in Figure 1B. For each selection round, 0.5 nmol RNA (∼3.0 x 10^14^ molecules) was refolded by heating to 90°C for 90 sec followed by cooling to room temperature for 5 min. This was followed by addition of 10X binding buffer to a final concentration of 1X (50 mM Tris-HCl [pH 7.5], 100 mM KCl, 50 mM NaCl, 1 mM MgCl_2_). For positive selection rounds against lattice or hexamer, 100 pmol of the respective protein was added (500 nM RNA to 100 nM CA protein). For negative selection rounds against hexamer, 2.5 nmol hexamer was added (500 nM RNA to 2500 nM hexamer). No protein was added to the binding reaction in the nitrocellulose trajectory. The final volume for each binding reaction was 1 mL.

Binding reactions were incubated at 37°C for 15 min followed by partitioning of the bound vs unbound RNA species. For positive selection rounds against lattice, partitioning was performed as described previously (*45*). Briefly, the binding reaction was centrifuged at 16,000 x g for 10 min at 4°C. Following centrifugation, the supernatant was removed, and the lattice pellet was washed with 1X binding buffer. The wash step was performed twice, followed by recovery of lattice-bound RNA using phenol/chloroform extraction and ethanol precipitation. For the rounds using hexamer or nitrocellulose, partitioning was performed using alkaline-treated nitrocellulose filters to decrease non-specific nucleic acid retention on the filter. Alkaline-treatment was performed by pre-incubating the filters in 0.5 M KOH for 20 min, washing them extensively with MilliQ water, and equilibrating them with 1X binding buffer for at least 45 min (*77*). During binding reaction incubation, the treated filters were placed on a sampling manifold and pre-wet with 1 mL 1X binding buffer. Immediately before applying the binding reactions, the filters were washed again with 1 mL 1X binding buffer under vacuum. The reaction was then applied and immediately washed with 1 mL 1X binding buffer. Filters remained under vacuum for an additional 2-3 min. For positive rounds using hexamer and rounds using nitrocellulose, RNA was recovered from filters by incubation with 400 μL extraction buffer (8 M urea, 10 mM EDTA, 50 mM NaCl), followed by phenol/chloroform extraction and ethanol precipitation. For negative rounds using hexamer, the RNA present in the flow-through (i.e., non-hexamer binders) was recovered using ethanol precipitation. Recovered RNA from all trajectories was reverse transcribed using ImProm-II Reverse Transcriptase (Promega, Madison, WI) and PCR amplified using Taq DNA polymerase for 10 cycles to generate the transcription template for the next round of selection.

### Illumina Sequencing

The libraries specified in Figure 1A were prepared for sequencing using a series of PCR steps to append the Illumina adapters and sequencing indices (Table S5) for the multiplexing of the libraries as previously described (*78*). Sequencing was performed on an Illumina NextSeq 500 (University of Missouri Genomics Technology Core) to generate 75 bp paired-end reads. Populations were demultiplexed and the paired-end reads were assembled into sequences using PANDASEQ (*79*) with a minimum overlap of 45 nt and a quality threshold of 0.6 to remove low-quality assemblies. Data preprocessing was performed using cutadapt (*80*) to trim the 5′ and 3′ constant regions from sequences and to discard any uncut sequences or sequences with lengths not within ± 6 nt of the expected size (56 nt) after trimming.

Processed populations were then analyzed using FASTAptamer software (*47*) and its R-based re-implementation FASTAptameR 2.0 (*48*) to count and normalize reads (FASTAptameR-Count), calculate fold enrichment and compare populations (FASTAptameR-Sequence Enrichment), group related sequences together (FASTAptameR-Cluster), analyze cluster composition (FASTAptameR-Cluster Diversity), and calculate fold enrichment of clusters (FASTAptameR-Cluster Enrichment).

Aptamers were named according to their cluster number and rank in L15 or (if the aptamer was not present in L15) their cluster number and rank in round L17+H7. For example, L15.6.1 was the seed sequence for cluster 6 within L15, while H7.10.1 was the seed sequence for cluster 10 in round L17+H7. Clusters were also named according to their cluster number in lattice R15 or (if the cluster was not one of the top 1000 clusters in lattice R15) their cluster number in round L17+H7.

### Enrichment Criteria for Identification of Aptamer Candidates

Aptamer sequences were identified as candidates for a binding screen using lattice and hexamer based on how they and their cluster of related sequences enriched and/or depleted in different selection trajectories.

For the original lattice selection (L8, L10 and L15), we focused on sequences that were the seed sequences of their respective clusters with a sequence RPM of at least 10 and a cluster RPM of at least 50 where both the sequence and cluster could not have depleted more than 2-fold (i.e., enrichment value greater than 0.5 from L8 to L10, L10 to L15, or L8 to L15).

Alternatively, if a sequence or cluster was not previously sampled in L8 or L10, such that an enrichment value could not be calculated, it was also considered.

For the nitrocellulose trajectory, candidates were not predicted to be hexamer binders if the sequence and its respective cluster had enriched greater than 5-fold in the nitrocellulose trajectory.

For the hexamer enrichment trajectory, some sequences had enriched (i.e., potential hexamer-binders) and others had depleted (i.e., potential lattice-specific binders) (Figure S4E-G). Four aptamers from the L15 library candidate list (L15.1.1, L15.2.1, L15.14.1, and L15.20.1) had enrichment values greater than 1 for both the individual sequence and its cluster from L15 to L17+H7 and from L17+H3 to L17+H7, suggesting that they could be capable of binding lattice and hexamer. Alternatively, sequences and their respective clusters absent in L15 but present in L17+H3 and L17+H7 were considered if they had a sequence RPM greater than 10 and a cluster RPM greater than 50 and had enriched more than 5-fold from L17+H3 to L17+H7. Only aptamer H7.10.1 met these criteria. Finally, for sequences and their respective clusters present only in L17+H7, sequences were considered if they had a sequence RPM greater than 10 and a cluster RPM greater than 50. Two aptamers (H7.12.1 and H7.20.1) met these criteria.

For the hexamer depletion trajectory, it was expected that any CA hexamer-binders would deplete, while lattice-specific binders would enrich or stay approximately the same.

Additionally, it was expected that significantly depleted sequences should be separately enriched in the lattice and hexamer trajectory. However, this prediction was observed only once, for L15.19.1, where the sequence had strongly depleted at both the sequence-and cluster-level and had enriched from L17+H3 to L17+H7 (Table 1).

**Table 1:**
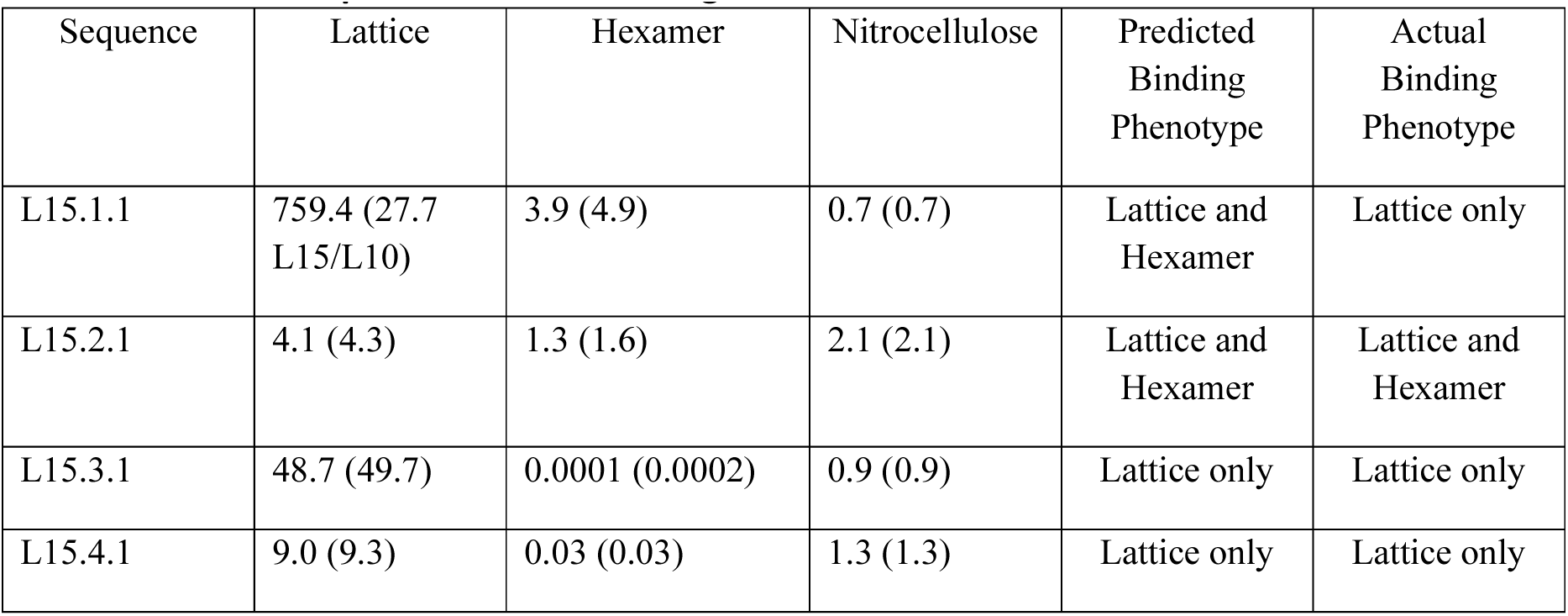
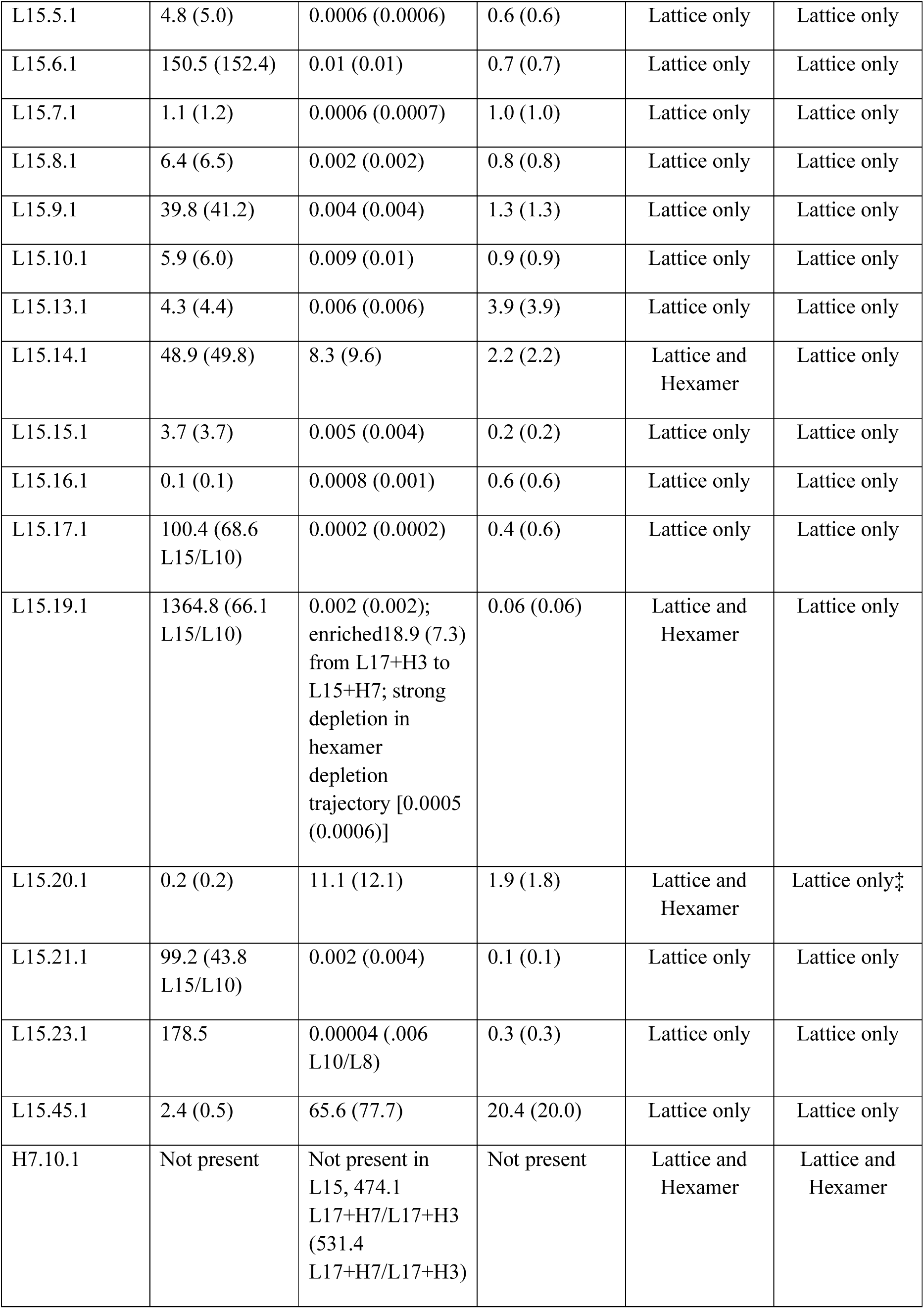
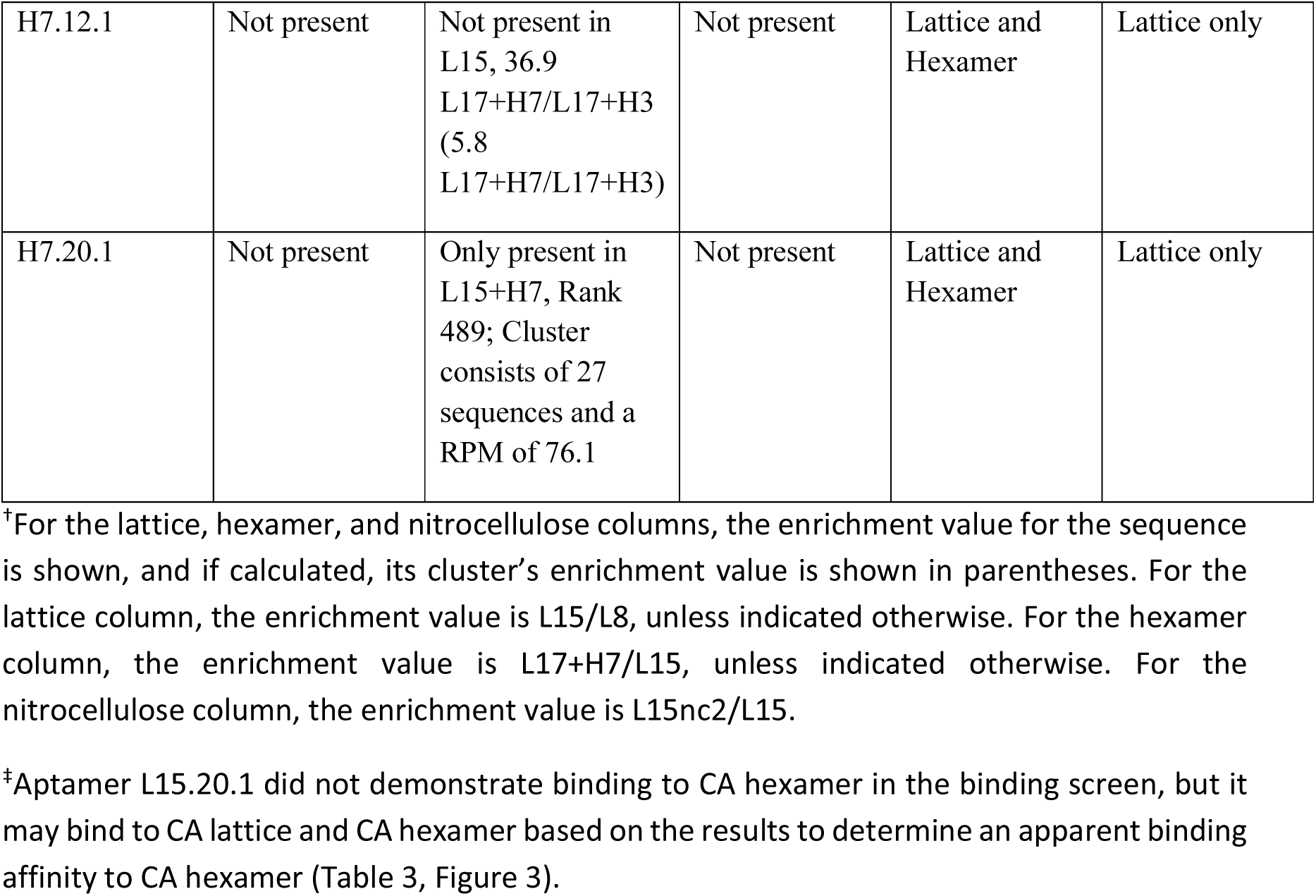
Candidate Aptamers for CA Binding Screen.

### DNA Templates and RNA Transcription

For each aptamer to be transcribed, DNA oligonucleotides were purchased as ‘left’ and ‘right’ halves from Integrated DNA Technologies (Coralville, IA). Oligonucleotides corresponding to the ‘right’ half were 5′-phosphorylated using T4 polynucleotide kinase (New England Biolabs, Ipswitch, MA). Equimolar amounts of ‘right’ strand, ‘left’ strand, and a complimentary bridge oligo were annealed and ligated using T4 DNA ligase (New England Biolabs, Ipswitch, MA). Ligated templates were PCR amplified using Pfu DNA polymerase, a forward primer to append the T7 promoter, and a reverse primer complimentary to the 3′ constant region. Amplified products were verified by size using agarose gel electrophoresis. The double-stranded DNA templates were then subjected to *in vitro* transcription and purified as described above.

### Nitrocellulose Filter Binding Assays

*In vitro* transcribed RNAs were treated with Antarctic phosphatase (Fermentas, Waltham, MA) to remove the 5’ terminal phosphate and then labeled by T4 polynucleotide kinase (New England Biolabs, Ipswitch, MA) in the presence of γ-^32^P-labeled ATP (PerkinElmer, Waltham, MA). Radiolabeled RNAs were gel-purified using denaturing PAGE as described above. To evaluate dose-dependent binding of aptamer libraries or individual aptamers to CA proteins, 50 nM 5′-radiolabeled and refolded RNA was incubated with varying concentrations of lattice, hexamer, or monomer in 1X binding buffer at 37°C for 15 min.

RNA-protein complexes were partitioned from unbound RNA by filtering samples through a pre-wet, alkaline-treated nitrocellulose filter under vacuum and immediately washing with 1 mL binding buffer. To determine radioactivity retained on the filter, filters were placed into scintillation vials with 4 mL Emulsifier-safe liquid scintillation fluid (Perkin Elmer, Waltham, MA) and analyzed using a liquid scintillation counter. A separate aliquot of the same RNA was incubated without CA protein and applied to filters under vacuum to determine non-specific nucleic acid retention. An unfiltered ‘No Wash’ sample was counted to determine the total amount of radioactivity present in each binding reaction. The fraction of RNA retained on the filter was calculated by dividing the radioactivity retained on the filter by the radioactivity present in the ‘No Wash’ sample. Three or more replicates were performed for each binding assay.

To determine the apparent dissociation constant (K_Dapp_) using nitrocellulose filter binding assays, approximately 20,000 counts-per-minute (cpm) of 5′-radiolabeled and refolded RNA was incubated with varying concentrations of lattice (0 nM lattice to determine background binding, or 25 to 10000 nM lattice) in 1X binding buffer at 37°C for 15 min. The nitrocellulose filter binding assay was then performed as described above. The background signal at 0 nM lattice was subtracted from all values in a dataset, and these values for the fraction of RNA retained on filter after background subtraction were fit to a 1:1 binding curve model (Y = B_max_*X/[K_D_+X]) using GraphPad Prism 6.2. In the equation, B_max_ is the maximum specific binding, K_D_ is the dissociation constant, X is the CA concentration (total monomer), and Y is the fraction of RNA retained on filter. Binding assays were performed at least in triplicate.

To determine the effect of buffer composition on aptamer binding, binding assays were performed as described above using either the standard 1X binding buffer or 1X binding buffer containing NaCl in place of KCl (50 mM Tris pH 7.5, 150 mM NaCl, 1 mM MgCl_2_). Binding assays were performed in triplicate.

### Structural motif search using MEME Suite

The top 1000 cluster seed sequences from L15 were searched for conserved structural motifs using the MEME discovery tool as part of MEME Suite(*49*). After identification of 20 motifs with a maximum width of 30 nucleotides, the sequence motifs were prioritized based on the number of sequences that contained each motif.

### Nitrocellulose Filter Binding Competition Assays

To determine whether an unlabeled competitor could compete with 5′-radiolabeled H7.10.1 for binding to the lattice, varying concentrations of unlabeled competitor (0.25-1 μM) were incubated with 1 μM lattice in 1X binding buffer at 37°C for 10 min. A separate sample with 0 nM competitor was prepared in parallel to determine the maximum binding of aptamer H7.10.1 within each assay. Unlabeled competitor RNA was prepared as described above. PF74 (Sigma Aldrich, St. Louis, MO) was prepared to a final concentration of 10% DMSO. 50 nM 5′-radiolabeled and refolded H7.10.1 was then added to the binding reaction, which was incubated at 37°C for 15 min. Nitrocellulose filter binding assays were performed and the fraction of radiolabeled H7.10.1 retained on the filter was calculated as described above. To calculate the relative fraction of radiolabeled H7.10.1 retained on filter, the fraction of radiolabeled H7.10.1 retained on filter at each concentration of competitor was divided by the fraction radiolabeled H7.10.1 retained on filter when no competitor was present (0 nM). These assays were performed in triplicate.

### Expression and Purification of the Stabilized Viral Cores

The method for the purification of the stabilized viral cores was adapted from Renner et al (*81*). HEK293FT cells were seeded in a 10 cm dish in DMEM medium containing 7.5% fetal bovine serum (FBS). When the cells reached 70-80% confluence, they were co-transfected using polyethylenimine (PEI; 1 mL of 1 mg/mL PEI per mg DNA) with 2.5 µg of pCMV-deltaR8.2 (A14C, E45C), 2.5 µg of HIV-1 packing plasmid encoding GFP (gift from Dr. Marc Johnson, University of Missouri), and 0.5 µg pMD-G encoding the VSV-G envelope. pCMV-deltaR8.2(A14C, E45C) was a gift from Wesley Sundquist (Addgene plasmid # 79047; http://n2t.net/addgene:79047; RRID:Addgene_79047Addgene) (*82*). pMD-G was a gift from Simon Davis (Addgene plasmid # 187440; http://n2t.net/addgene:187440; RRID:Addgene_187440) (*83*). After 48 hr, supernatants containing VSV-G-pseudotyped viruses were collected, centrifuged for 5 min at 1000 x g to remove cellular debris, and filtered through 0.45 µM filters. The filtered viral supernatant was then overlaid onto a density gradient (2 mL sterile 20% sucrose in 1X PBS and 1 mL of 5% sucrose in 1X PBS containing 2% Triton X-100) in Beckman Coulter Polypropylene tubes. The stabilized viral particles were then centrifuged at 30,000 RPM for 16 hr at 4°C using a Beckman Coulter Optima L-90K ultracentrifuge. After ultracentrifugation, 0.5 ml fractions were removed carefully without disturbing the density gradient. The capsid pellet at the bottom of the tube was then washed with 1X PBS and centrifuged at 30,000 RPM for 1 hr. After ultracentrifugation, the stabilized viral cores were resuspended in 1X PBS. To assess the integrity of the purified capsid cores, the fractions of the density gradient ultracentrifugation and pelleted viral core were analyzed through SDS-PAGE followed by western blot (Figure S7). Briefly, after protein transfer to a PVDF membrane, the membrane was blocked with 5% milk and washed three times. The membrane was incubated with anti-p24 monoclonal antibody at 1:2000 dilution overnight, washed, and then incubated with anti-mouse HRP-labeled secondary antibody at 1:4000 dilution. After three washes, the blot was developed using SuperSignal^TM^ West Pico PLUS chemiluminescence substrate (ThermoFisher Scientific, Waltham, MA) using Odyssey® Fc Imaging System (LI-COR Biosciences, Lincoln, NE).

### Enzymatic Probing of Aptamer Structures

The 5′ radiolabeled aptamer structures were probed using enzymatic digestions (*57*). To generate the OH ladder, 1 μL 5′ radiolabeled RNA (50,000-150,000 cpm/μL) was incubated in 50 mM sodium carbonate pH 9.0 buffer at 90°C for 10 min. The alkaline digestion reaction was stopped by adding an equal volume of colorless loading dye (10 M urea, 15 mM EDTA) and snap cooling in a bath of dry ice and ethanol. A denaturing T1 digestion was performed by incubating 1 μL of 5′ radiolabeled RNA (50,000-150,000 cpm/μL) in 25 mM sodium citrate pH 5.0 buffer containing 7 M urea and 10.5 mM EDTA with 1 U RNase T1 (ThermoFisher Scientific, Waltham, MA) at 55°C for 5 min. The reaction was stopped by adding an equal volume of colorless loading dye and snap cooling. For the native RNase T1, S1 nuclease, and VI RNase digestions, the 5′ radiolabeled RNA (50,000-150,000 cpm per reaction) was first heated in water at 90°C for 5 min and then allowed to cool at 0.1°C/sec in a thermocycler.

When the temperature reached approximately 65°C, 10X capsid binding buffer was added to the RNA to a concentration of ∼1.1X. Once at room temperature, the RNAs were placed on ice. Prior to adding an enzyme, reactions were equilibrated at 37°C for 2 min. For the native T1 digestion, 0.1 U was added to the reaction and incubated at 37°C for 2 min. For S1 nuclease (ThermoFisher Scientific, Waltham, MA), reactions were performed with and without the addition of 1 mM ZnSO_4_ prior to equilibration at 37°C. 20 U of S1 nuclease were added to the reaction and incubated at 37°C for 5 min. For RNase VI (Ambion, Austin, TX), 0.005 U were added to the reaction and incubated at 37°C for 5 min. Enzymatic digestions were stopped by adding an equal volume of colorless loading dye and snap cooling. Reactions were run on a 0.4 mm thick, 8% denaturing polyacrylamide gel (TBE-8 M urea). The gel was exposed to a phosphoimager screen overnight at -20°C and the phosphoimager screen was scanned using the Typhoon FLA 9000 phosphoimager (GE Healthcare, Chicago, IL). Experiments were performed in triplicate to establish reproducibility. Representative gels are shown.

### Generation of Covariance Models using Infernal

Covariance models were generated following a protocol similar to that described previously for HIV-1 RT aptamers (*78*). For the top 10 clusters and cluster 20 from L15, the 300 most abundant sequences within each cluster were aligned using MAFFT (*59*) and these alignments were used to predict a secondary structure using RNAalifold (weight of the covariance term = 0.6, penalty for non-compatible sequences = 6) (*84*). Both the MAFFT alignment and RNAalifold prediction were performed within the Jalview Desktop application (*85*). The results from RNAalifold were exported as a Stockholm file within Jalview. To use the Stockholm file in Infernal, the centroid structure was renamed SS_cons, and an extra ‘-’ was removed from the end of the centroid structure line. The input Stockholm file containing the alignment and centroid structure predicted by RNAalifold were then used to generate a covariance model (CM) using the Infernal program (*60*). Infernal was then used to calibrate and search the covariance models against the population. CMs were searched against the top 1000 cluster seed sequences from L15. For all CM searches, seed sequences matched with the CM if the expectation score was below an expectation score cut-off of 0.1. The default value is 0.01, which means about one false positive would be expected every 100 queries on average. To increase the number of hits, an expectation cut-off score of 0.1 was used, as done previously (*78*). Groups of clusters that were identified from the CM search were then aligned to the CM and used to generate a new CM. This process was repeated until either no more sequences matched the CM or up to 10 rounds total. The results from the final CM search were then visualized using R2R (*86*) to visualize sequence and/or structure conservation.

### Microscale Thermophoresis (MST)

Binding affinities of aptamer or the arbitrary RNA control for lattice were determined by measuring thermophoresis of RNAs containing a 3′ tail extension annealed to a Cy5-labeled anti-tail oligonucleotide in the presence of increasing concentrations of protein. Reaction mixtures containing 25 nM RNA-tail/Cy5-anti-tail complexes and increasing concentrations of CA proteins (39–5000 nM) were prepared in 1X binding buffer containing 0.5% pluronic acid. Reaction mixtures were incubated for 15 min at 37°C and then loaded into capillaries. Thermophoresis was monitored on a Monolith NT.115 MST instrument (NanoTemper Technologies GmbH, Munich, Germany) at 20% LED power and medium MST power with 20 sec MST-on time, and data were analyzed with MO.Affinity software (version 2.3) using a 1:1 binding model (Nano Temper Technologies, CA). To normalize the replicates, ΔF norm values were calculated by subtracting the F norm value from 39 nM at all concentrations. The final ΔF norm data were plotted using GraphPad Prism (GraphPad Inc., La Jolla, CA). Experiments were performed four times.

### RNA Mutational Profiling (MaP)

RNA (50 nM) was refolded as described above. 25 µl of folded aptamer RNA was mixed with 12 µl of 1 M bicine (pH 8.3 at 25°C) and incubated at 37°C for 5 minutes. Next, 4 µl of 1 M TMO (trimethyloxonium tetrafluoroborate) solution, prepared in 1:2 (v/v) nitromethane/sulfolane (*58*), was added to the RNA solution and incubated at 37°C for 10 seconds. The reaction was quenched by adding 200 µl of stop solution (300 mM Tris, 300 mM NaCl, 0.5% SDS, 2 mM EDTA, pH 7.4) containing carrier RNA (3 µg/ml), followed by ethanol precipitation. Half of the dissolved ethanol-precipitated RNA (10 µl) was mixed with 0.2 µM of 56N Reverse Primer (5’-TATGGACTTACCTACCTTATGCCC-3’) and incubated at 95°C for 15 seconds, 65°C for 5 minutes, followed by 4°C for 2 minutes. Then, 8 µl of 2.5 x MaP buffer (125 mM Tris, 187.5 mM KCl, 15 mM MnCl_2_ 25 mM DTT and 1.25 mM dNTPs, pH 8.0) (*87*) was added, and the combined solution was incubated at 42°C for 2 minutes. Finally, 1 µl of M-MLV Reverse Transcriptase (Promega) was added, and the reaction was performed at 42°C for 3 hours. The resulting cDNA was purified using the NEB Monarch DNA Cleanup Kit (NEB, T1030) and eluted with 10 µl of elution buffer.

Sequencing libraries were prepared from the cDNA using nested PCR to incorporate Illumina-TruSeq DNA and RNA UD Indexes and adaptors (IDT) for each sample using Library primers and UDI primers. The final amplicons of each sample were assessed by agarose gel and purified using a NEB DNA silica column (NEB, T1030). The pooled library was sequenced using the 2 x 150 paired-end protocol on the Illumina HiSeq platform (Novagene). Data were analyzed using ShapeMapper 2 (*88*).

### Circular Dichroism and UV-Visible Spectrophotometry

Circular Dichroism (CD) measurements were collected using a Jasco J-1500 spectrophotometer with an MPTC-511 Peltier 6-position cell holder from 0°C to 95°C, with a 3°C/min ramp, and measurements taken every 5°C. Four accumulations were collected for all spectra from 340-200 nm at a scanning speed of 200 nm/min. Absorbance and High Tension (HT) voltage measurements were collected simultaneously. Initial absorbance values of ∼0.8 were collected at room temperature and verified with a Jasco V-750 spectrophotometer. Further, absorbance values were monitored at all temperatures to account for hyperchromicity shifts, with values reaching no higher than ∼1.2 within regions of interest. The HT voltage remained below 500 V for all collected spectra between 340-200 nm. All aptamers were precipitated and dissolved in nuclease-free water, being stored at - 20°C prior to use. Aptamers were denatured at 95°C for 1 min and placed in their respective folding buffers at concentrations of 10 µM prior to the collection of any spectra. The quartz cuvettes employed had pathlengths of 0.1 cm and required a cuvette adapter for use in both spectrophotometers. Standard curve data for each aptamer was collected using a NanoDrop One microvolume spectrophotometer and found to be in agreement with the experimental data. Spectral analyses, Lagrange interpolation, and Van’t Hoff thermodynamic calculations were performed using a combination of Jasco Spectra Analysis software, Microsoft Excel, and GraphPad Prism.

### Folding Calculations

Alpha (*α*) values were calculated from ellipticity data collected during CD/UV-Vis thermal denaturation studies of each aptamer under variable cationic conditions. This data was plotted and interpolated, allowing for determination of melting temperatures (Tm), where *α* = 0.5, and facilitated examination of the overall fold at a given temperature, where (*89*):

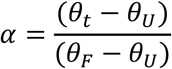

Derivative plots were also created to verify the fit with calculated Tm values (*90, 91*). Folding equilibrium constant (*K*) values were calculated to determine the free energy (*ΔG*) of folding for each aptamer under each condition (*89, 92*):

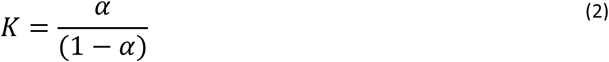

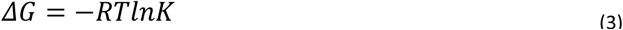

Finally, Van’t Hoff plots and calculations were performed to further clarify the thermodynamic properties of each aptamer (*89, 92*):

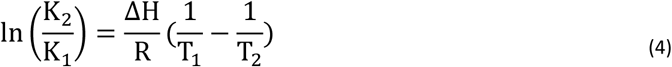

### Nuclear Magnetic Resonance Spectroscopy

Analyses of the ^1^H imino proton regions of each aptamer were performed using a Bruker Advance III 800 MHz Spectrometer with a TCI cryoprobe and Sample Xpress changer. Concentrations ranged from ∼100-150 µM, and were dissolved in 90% H_2_O and 10% D_2_O, with folding buffer conditions consisting of 10 mM Tris, 150 mM KCl, and 1 mM MgCl_2_ at a pH of 6.3. Applied temperatures varied from 10-35°C to examine potential structural changes. Data treatment and analyses were performed using NMRViewJ and Bruker TopSpin software.

### Lattice Pulldown

Indicated aptamer RNAs were first annealed to a biotinylated oligonucleotide complementary to the 3′ constant region. The complexes were then incubated with streptavidin magnetic beads. Cellular lysates were harvested from HEK293FT cells and spiked with assembled lattice protein. Aptamer-bead complexes were then incubated with the lattice spiked lysates. Complexes were separated from the lysates using a magnet, washed thoroughly, and eluted from the beads using heat and buffer containing excess biotin. Eluates were then subjected to denaturing SDS-PAGE and western blot. The membrane was incubated with anti-p24 monoclonal antibody at 1:2000 dilution overnight, washed, and then incubated with anti-mouse HRP-labeled secondary antibody at 1:4000 dilution. After three washes, the blot was developed using SuperSignal^TM^ West Pico PLUS chemiluminescence substrate (ThermoFisher Scientific, Waltham, MA) using Odyssey® Fc Imaging System (LI-COR Biosciences, Lincoln, NE).

### Statistical Analysis

To determine whether there was a statistically significant difference between two sets of samples, P values were calculated using an unpaired t test with Welch’s correction computed by GraphPad Prism.

### Structure Prediction Software

Structure prediction throughout this work was performed using Mfold (*71*) and the NUPACK web server (www.nupack.org) (*72*). Prediction of potential G-quadruplex structures was performed using QGRS Mapper (https://bioinformatics.ramapo.edu/QGRS/index.php) (*67*).

## RESULTS

### HIV-1 Capsid Nomenclature

The experiments detailed here utilize various HIV-1 capsid assembly forms. For the purposes of this study and throughout the remainder of the text, we will utilize the following nomenclature: capsid protein (CA), A14C/E45C CA hexamer lattice tubes (lattice), A14C/E45C/M184A/M185A soluble CA hexamer (hexamer), wildtype soluble CA monomer (monomer), and A14C/E45C mature capsid core (stabilized mature capsid core).

### CA differentiation selection

We previously described selection of aptamers using lattice as a target (*45*). The selected aptamer libraries exhibited dose-dependent binding to lattice but not to hexamer or monomer (*45*), suggesting that the majority of aptamer sequences present within the library were lattice-specific binders. As the lattice contains repeated units of lower order CA assembly forms (e.g. monomer and hexamer), we reasoned that the lattice round 15 (L15) library may also contain sequences capable of binding these forms (Figure 1A).To identify aptamer subsets from within the L15 library that were either specific for lattice or capable of binding both lattice and hexamer, we performed a differentiation selection using L15 as the starting point, with lattice and hexamer as the protein targets (Figure 1B). The lattice used here for aptamer selection is composed of cross-linked A14C/E45C CA hexamers that retain higher-order assembly under physiological conditions, while hexamer is composed of cross-linked A14C/E45C/W184A/M185A CA hexamers that are incapable of forming higher-order CA assemblies under physiological conditions (*46*).

The CA differentiation selection included three selection trajectories (Figure 1B). To identify sequences capable of binding both lattice and hexamer (trajectory 1), we performed four enrichment rounds that toggled between hexamer and lattice, followed by five enrichment rounds using hexamer alone. Partitioning of bound from unbound sequences was performed using centrifugation for lattice (*45*) and nitrocellulose filter retention for hexamer. In trajectory 1, sequences that bound either target were retained, while sequences that were unable to bind either target were discarded. To identify lattice-specific binders, trajectory 2 incorporated depletion for sequences that bind hexamer. Depletion rounds retained sequences that did not bind hexamer (flow-through), and these rounds were toggled with enrichment rounds to maintain lattice binding. Importantly, hexamer protein concentration for each depletion round was in excess of the RNA aptamer library concentration. Because retention on nitrocellulose filters was used to partition sequences that bound hexamer from those that did not, trajectory 3 sought to identify sequences that non-specifically bound nitrocellulose. Binding reactions for trajectory 3 were prepared with RNA aptamer libraries in the absence of protein and applied to nitrocellulose filters under vacuum. Sequences that remained on the filter after washing were retained.

Aptamer libraries generated from the lattice and differentiation selections were next subjected to high throughput sequencing (HTS) analysis and named according to the number of partition steps against the different selection targets and the type of selection step performed (Figure 1B, within circles). More than 34 million high-quality reads remained following data preprocessing to merge paired end reads, trim off constant regions, and remove low quality or ambiguous reads. Processed libraries were analyzed using the FASTAptamer toolkit (*47*) and its R-based reimplementation, FASTAptameR 2.0 (*48*). Read counts and the number of unique aptamer sequences for each library are shown in Table S1. Comparison of the percentage of unique sequences present within the aptamer libraries (unique sequences/total reads) demonstrated that L15 and the subsequent differentiation selection rounds were substantially more converged than L8 and L10 (Table S1 and Figure S1). Most aptamer reads had a sequence length of 56 nucleotides (nt) (Figure S2) with minimal insertions or deletions during amplification steps (i.e., PCR and *in vitro* transcription). Sequences were clustered into families of closely-related sequences using FASTAptameR-Cluster, with Levenshtein edit distance (LED) set to 6. The top one thousand clusters were generated for each library, except for L8 where the top one hundred clusters were generated due to the large number of unique sequences present in the library.

### Identification of aptamer candidates and binding validation

Aptamer sequences were identified as candidates for a binding screen based on how they and their cluster of related sequences enriched and/or depleted in different selection trajectories. Pairwise comparisons between libraries were performed using FASTAptameR-Sequence Enrichment to determine the enrichment values of individual sequences, defined as the ratio of (RPM in round y [RPM_y_]) to (RPM in round x [RPM_x_]). While enrichment values can only be calculated for sequences present in both libraries being compared, many sequences were present only in one or the other (Figure S3). Plotting normalized read frequencies (RPMs) for sequences present in each library being compared (Figure S4) identified sequences that enriched or depleted (above or below the y = x line, respectively) and sequences whose frequencies stayed approximately the same had points that fell on or near the *y* = *x* line.

Because the original L15 library was used as the branch point for the differentiation selection, we expected sequences identified as candidates to bind lattice. Sequences that depleted in the lattice and hexamer trajectory but did not significantly deplete in the hexamer depletion trajectory were predicted to be lattice-specific binders. In contrast, sequences that enriched in the lattice and hexamer trajectory and significantly depleted in the hexamer depletion trajectory were predicted to be capable of binding both lattice and hexamer. Those predicted to bind both lattice and hexamer were chosen for further study if they did not also significantly enrich (≥5-fold) in the nitrocellulose trajectory, in which case they were considered false positive hits.

Table 1 lists 23 aptamers identified informatically for screening, their enrichment values from different selection trajectories, and their predicted binding phenotype based on their enrichment or depletion profiles. Aptamer binding to lattice and hexamer was determined experimentally using nitrocellulose filter binding assays. As expected, all 23 aptamers exhibited dose-dependent binding to lattice, while two aptamers exhibited clear binding to hexamer, H7.10.1 and L15.2.1 (Figure 2, Table S2). Aptamers H7.10.1 and L15.2.1 exhibited dose-dependent binding to hexamer compared to an arbitrary control (Arb); however, L15.2.1 also exhibited more non-specific binding to nitrocellulose (Figure 2B, Table S2).

**Figure 2:**
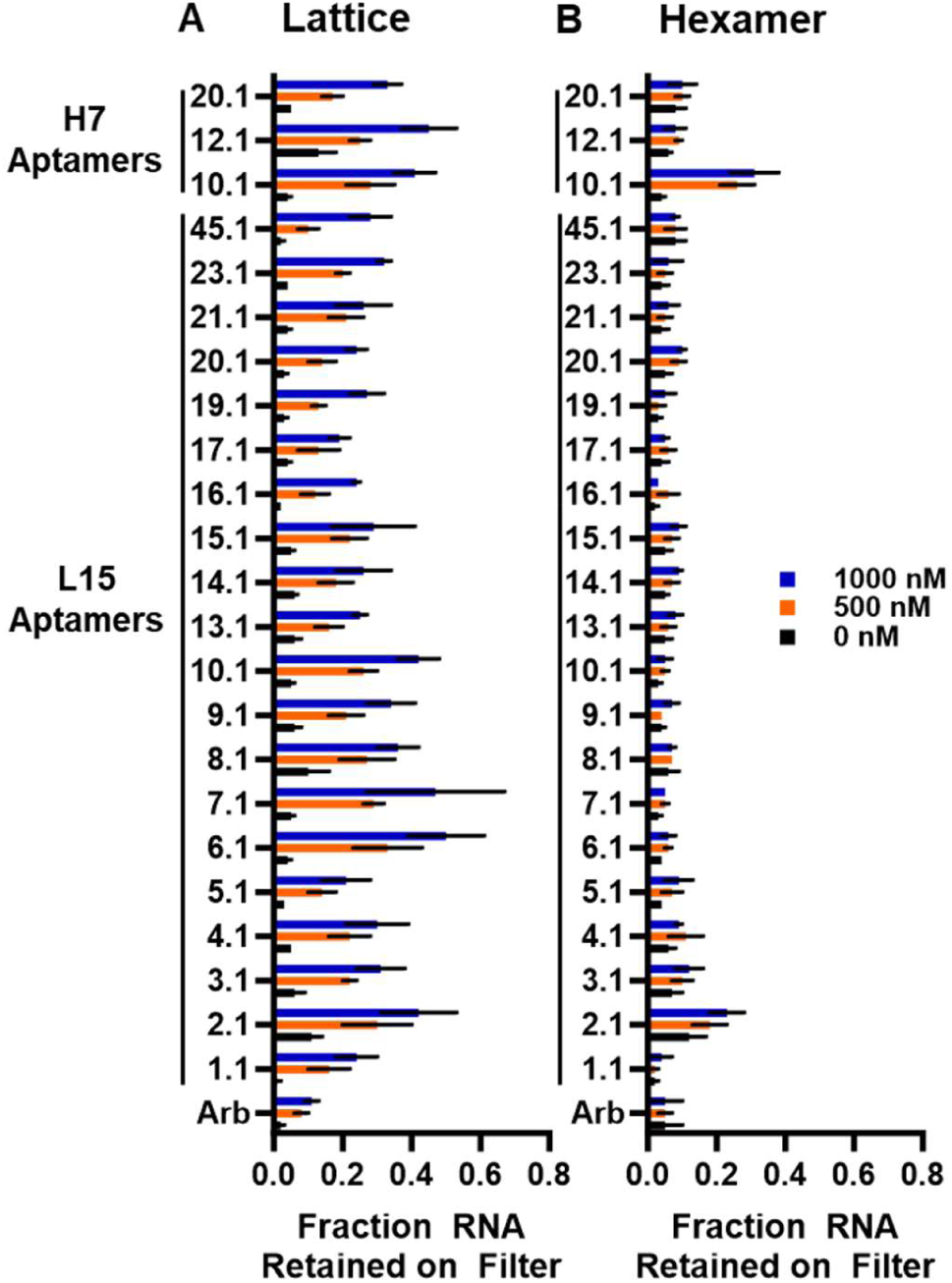
CA binding by candidate aptamers. The binding of candidate aptamers to CA lattice (A) and CA hexamer (B) was assessed using nitrocellulose filter binding assays. Experiments were performed in triplicate at minimum. Values plotted are the mean ± SD. Results from statistical tests are in Table S2.

Other aptamers demonstrated slight dose-dependent binding to hexamer (e.g., L15.5.1 and L15.20.1), but not to the degree of L15.2.1 and H7.10.1. Importantly, our analysis identified subsets of aptamers that were lattice-specific or that were capable of binding both lattice and hexamer, with both binding patterns emerging from an already highly converged pre-enriched aptamer library that did not contain many sequences capable of binding hexamer.

### Identification of multiple CA-binding sequence families

Three widely recurring sequence motifs (Figure S5) were identified by analyzing the seed sequences from the top 1000 clusters in the L15 population using MEME Suite (*49*). Table 2 lists the top three most abundant sequence motifs, the number of seed sequences in which they were present, and the candidate aptamers from the prioritized list in Table 1 that contain the motifs. The top motifs included a highly G-rich motif (motif 1), a highly conserved GUGUAU motif (motif 2), and a conserved ACCUCC motif followed by a G-rich region (motif 3). Of our aptamer candidates, ten contained motif 1, six contained motif 2, and three contained motif 3. We chose aptamer L15.7.1 to represent motif 1 and aptamer L15.6.1 to represent motif 2, as they both demonstrated significant binding to lattice and no dose-dependent binding to hexamer. Aptamer L15.20.1 was chosen to represent motif 3, as it had been initially predicted to bind both lattice and hexamer and demonstrated slight dose-dependent binding to hexamer in our screen, suggesting some affinity for hexamer.

**Table 2:**
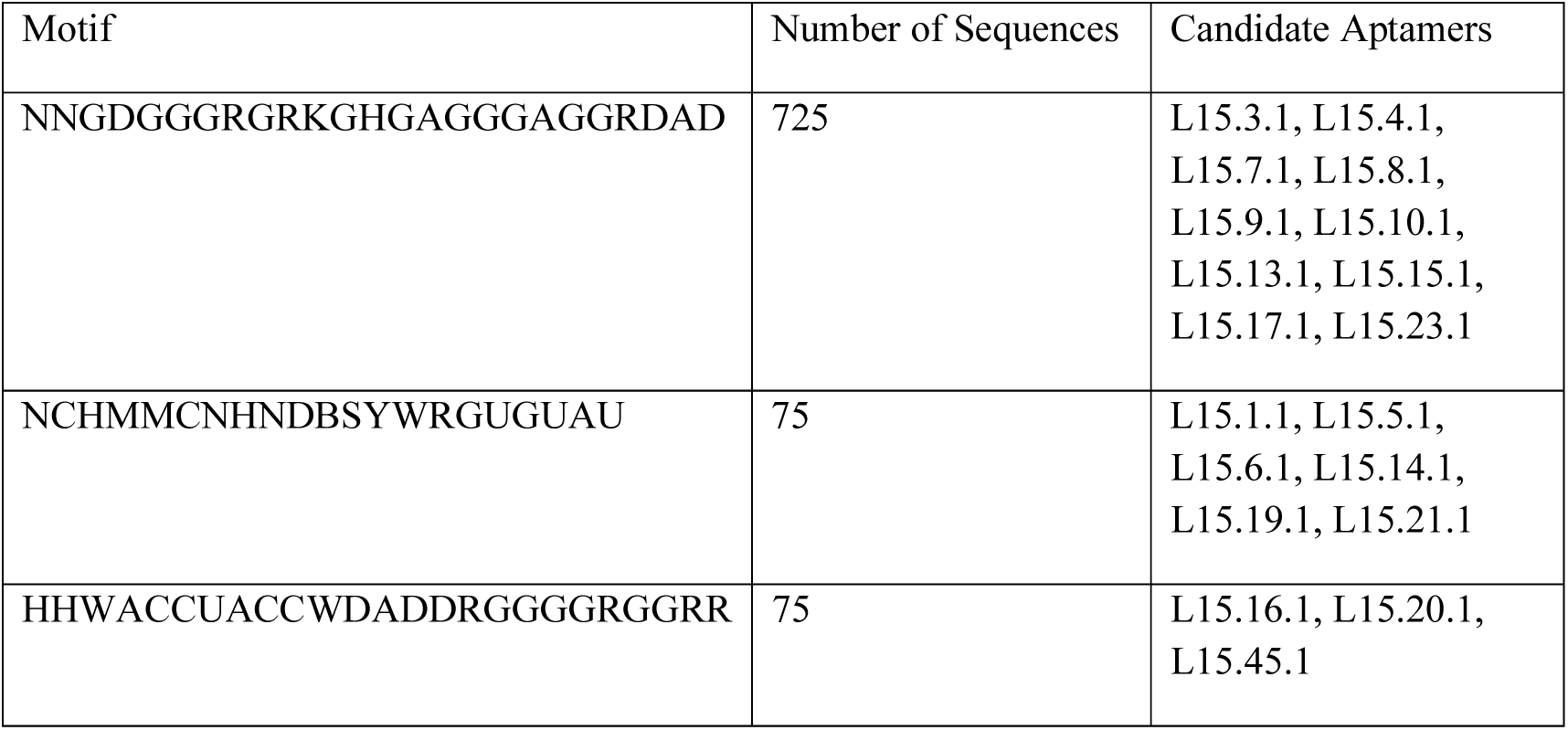
Top Three Most Abundant Sequence Motifs Identified in the Lattice Round 15 Library.

We also prioritized aptamer H7.10.1, one of the only lattice and hexamer binders identified. Somewhat unsurprisingly given its distinct recognition pattern, aptamer H7.10.1 did not contain any of the motifs identified from the L15 top 1000 cluster seed sequences, nor did it contain any of the twenty most abundant sequence motifs identified from L17+H7, suggesting that it may contain unique sequence and/or structural elements.

### Candidate aptamer binding to CA assembly forms

Apparent binding affinities (Kd_app_)for the prioritized aptamers were determined for both lattice (Table 3, Figures 3A and S6) and hexamer (Table 3, Figure 3B). All four aptamers bound lattice and aptamers H7.10.1 also bound hexamer, with Kd_app_ ranging from low nM to μM depending on the assay. Aptamer L15.20.1, which was originally predicted to bind both lattice and hexamer but did not demonstrate significant binding to hexamer in the initial binding screen (Table 1 and Figure 2B), bound hexamer with a Kd_app_ of 790±400 nM (Table 3, Figure 3B). Interestingly, while the Kd_app_ of L15.20.1 for hexamer was similar to its Kd_app_ for lattice in MST assays (Table 3), we observed substantial differences in Kd_app_ for L15.20.1 binding to lattice in nitrocellulose filter binding (4400±1600 nM) versus microscale thermophoresis (MST; 730±390 nM) assays.

**Figure 3:**
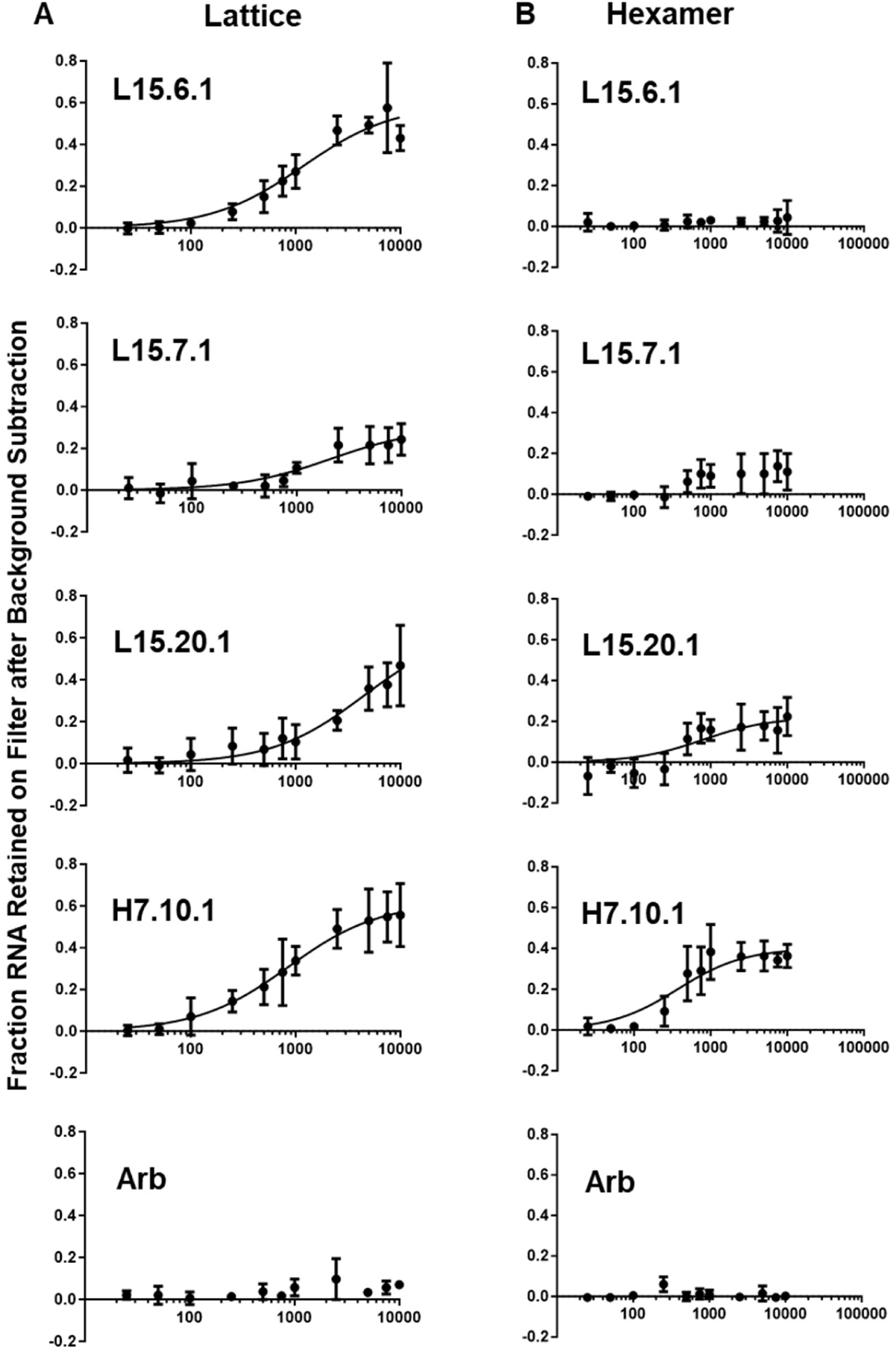
Apparent Kd Binding Curves for Representative Aptamers to Lattice and Hexamer. Binding curves were generated from nitrocellulose filter binding assays (n = 3). Radiolabeled RNAs were incubated in the presence of increasing concentrations of lattice (A) or hexamer (B), as determined on the basis of CA monomer, and applied to nitrocellulose filters. The fraction of RNA retained on the filter after background subtraction is plotted ±SD.

**Table 3:** Apparent Binding Affinities (K_D_) of Aptamers and Arbitrary Sequence to CA Lattice and CA Hexamer.

| RNA | CA Lattice $K_D$ (nM, nitrocellulose filter binding assay) | CA Lattice $K_D$ (nM, microscale thermophoresis) | CA Hexamer $K_D$ (nM) |
| --- | --- | --- | --- |
| L15.6.1 | 1181 ± 260 | 655 ± 279 | Did not bind |
| L15.7.1 | 2122 ± 823 | 1176 ± 336 | Did not bind |
| L15.20.1 | 4413 ± 1635 | 733 ± 395 | 793 ± 403† |
| H7.10.1 | 856 ± 166 | 705 ± 230 | 372 ± 104 |
| Arb | Did not bind | Did not bind | Did not bind |

†Aptamer L15.20.1 may bind to CA hexamer but it had high non-specific binding to the nitrocellulose filters.

We next tested aptamer binding to monomer, an assembly form present in both lattice and hexamer that also offers potential binding sites, such as the cyclophilin A binding loop. Aptamers L15.7.1, L15.20.1 and H7.10.1 were unable to bind monomer (Figure 4A), suggesting their binding interface is present or accessible only on hexamer and/or lattice assembly forms. These results extend similar observations from our prior study, where we showed that aptamer L15.6.1 (CA15-2) was also unable to bind monomer (*45*).

**Figure 4:**
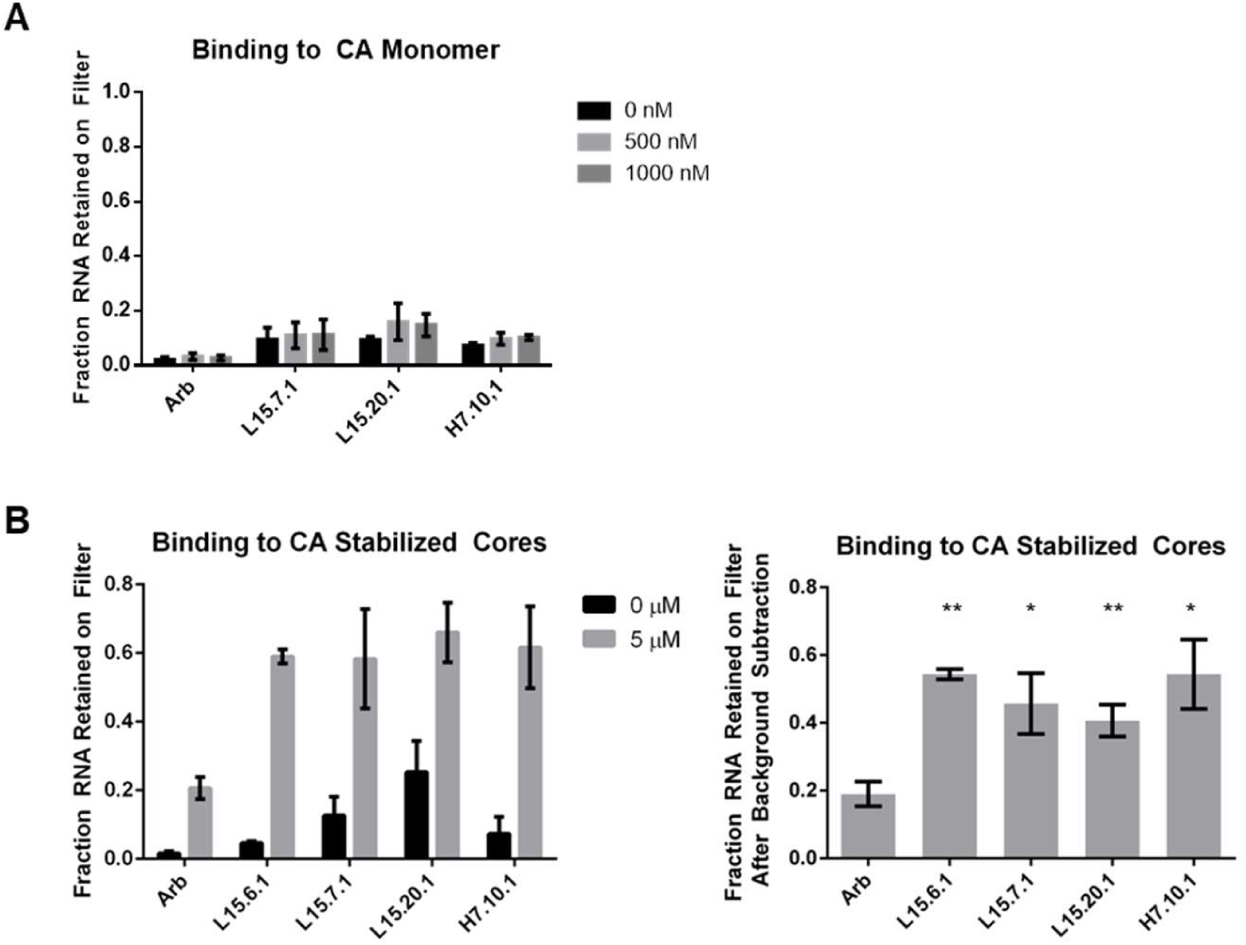
Binding of representative aptamers to CA monomer and stabilized CA cores. Binding to CA monomer (A, n = 4) and stabilized capsid cores purified from viral particles (B, n = 3) was accessed using nitrocellulose filter binding assays. (B, Right) The background binding to nitrocellulose was subtracted for each replicate to directly compare specific RNA binding to cores. Statistical comparisons were made for each aptamer versus the magnitude of binding of the arbitrary control. * (P < 0.05), ** (P < 0.01). Values plotted are the mean ± SD.

A long-term goal of these investigations is to develop aptamers as tools to study CA assembly forms in cells; thus, we determined whether our aptamers could bind a biologically relevant form of CA. During HIV-1 infection, the hexamer-hexamer interactions found in lattice are present in the mature capsid core, along with additional pentamer units at sites of curvature (*50*). To determine whether the aptamers could bind the mature capsid core, we isolated stabilized mature capsid cores from viruses (Figure S7) and evaluated binding. We observed that aptamer binding to the stabilized cores was significantly greater than binding by Arb (Figure 4B), suggesting that the aptamers could potentially interact with the mature capsid core in cells.

Higher order CA assembly forms contain known binding pockets for host factors and small molecules (*51–53*), such as the PF74 binding site located at the NTD (N-terminal domain)- CTD (C-terminal domain) interface of two monomers within assembled hexamers (*54*). Because aptamer H7.10.1 was found to bind both lattice and hexamer, we evaluated the ability of H7.10.1 to compete for binding with PF74. Unlabeled competitor (250-1000 nM) was first allowed to bind lattice (1 μM total monomer), followed by the addition of 50 nM radiolabeled H7.10.1. As expected, unlabeled aptamer H7.10.1 competed with itself for binding to lattice. However, similar to our prior results with lattice-specific aptamer L15.6.1 (*45*), PF74 did not compete with H7.10.1 (Figure S8), suggesting that the binding sites for PF74 and H7.10.1 are non-overlapping or that binding of PF74 does not block the binding of H7.10.1. Of note, the Kd of PF74 for hexamer is 262 nM (*54*), while H7.10.1 binds with a Kd_app_ of 372 ± 104 nM (Table 3, Figure 3B).

### Sequence and/or structural requirements for aptamer binding to lattice

In the absence of ligand, RNA often exists as an ensemble of conformations, presenting a significant challenge for the determination of apo RNA structure (*55, 56*). Despite this challenge, the study of apo aptamer RNA structures is of critical importance to advance our understanding of the binding mechanisms for aptamers to facilitate their development as molecular tools (e.g. affinity reagents) (*56*). Thus, we used a combination of different methods to probe the sequence and structural requirements for aptamer binding to lattice, as detailed below.

*Aptamer L15.6.1*: We previously evaluated truncations and modifications for L15.6.1, including annealing of complementary oligos to the 5′ and 3′ constant regions (*45*). We also predicted formation of a stem-loop structure consisting of nucleotides 30 through 60, with the identified GUGUAU motif (Figure 5A; nucleotides 43-48) present within a loop (*45*). To test the predicted secondary structure, we first performed nuclease digestion under native conditions using T1 RNase (cleaves after accessible G residues), S1 nuclease (cleaves single-stranded regions), and V1 RNase (cleaves double-stranded regions) (Figures 5A and S9). The S1 reactions were done in the absence and presence of 1 mM ZnSO_4_ because the S1 nuclease has optimal activity in the presence of 1 mM Zn^2+^(*57*). Cleavage patterns from the nuclease digestions were largely in agreement with the predicted secondary structure (Figure 5A and S9). Similarly, mutational profiling (MaP) performed using trimethyloxonium (TMO) in bicine buffer, which modifies accessible nucleotides in single-stranded regions (*58*), identified regions of high TMO reactivity in the predicted loops and single-stranded regions (Figure 5B). A covariation model (CM) generated using MAFFT (*59*) and Infernal (*60*) also positioned the GUGUAU motif within a loop (Figure 5C). Collectively, these data supported the predicted secondary structure.

**Figure 5:**
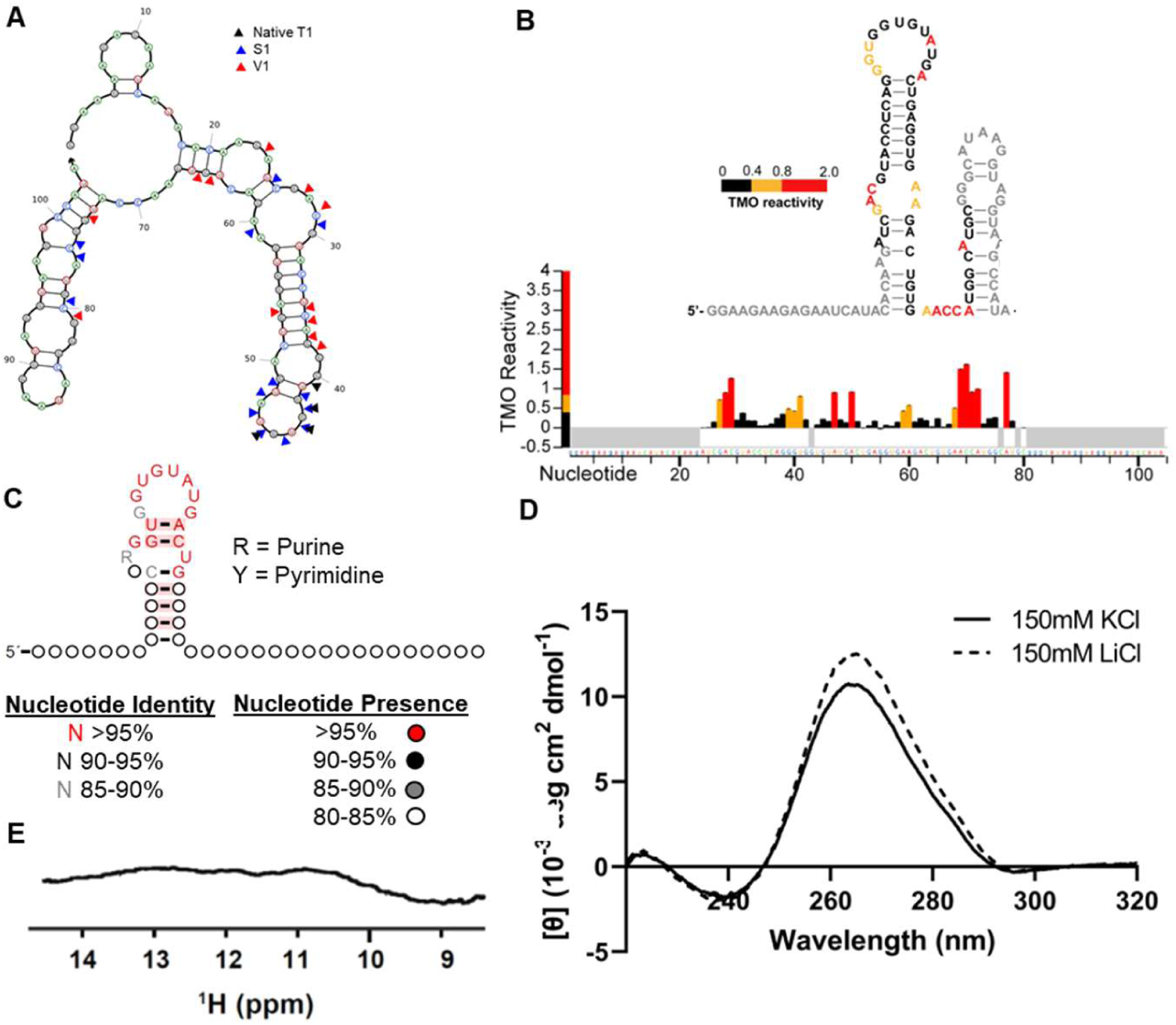
Exploration of the secondary structure of aptamer L15.6.1. (A) Mapping the cleavage products onto the predicted secondary structure of L15.6.1. Accompanying representative gel in Figure S9. (B) Regions of TMO modification in L15.6.1 MaP experiment. Top: Predicted secondary structure of L15.6.1 where the nucleotides are colored from least (black) to most (red) TMO reactive. Nucleotides in the constant regions that were not mapped by TMO modification are colored gray. Bottom: The magnitude of TMO reactivity is shown at each nucleotide position. (C) Covariance model generated starting from the cluster of L15.6.1. Red shading of base pairs denotes no variation in the base pair when present. (D) CD spectra of L15.6.1 in 150 mM KCl or LiCl. (E) NMR spectrum of L15.6.1.

While the data above supported the predicted secondary structure, we noted that our aptamer sequences, including L15.6.1, are G-rich (>30% G). G-rich motifs in RNA, such as G-quadruplex-like structures, tend to be particularly tight and stable (*61, 62*), which could benefit downstream development of these aptamers as research tools. As recent evidence supports that in RNA, G-quadruplexes can adopt non-traditional structures that may be comprised of more than just successive guanine nucleotides (*63–65*), we sought to expand our structural analysis to examine potential formation of G-quadruplexes and related structures (referred to from here as G4s) using circular dichroism (CD), and 1D ^1^H-NMR.

Changes in the ellipticity maxima at ∼265 nm and minima at ∼240 nm between RNA CD spectra under different monovalent cation conditions are known to occur with aptamers that contain G4s (*66*). Further, examination of the imino region in spectra generated using 1D ^1^H-NMR can identify uncommon G-rich motifs, including potential G-tetrads and/or non-canonical base-pairing. CD analysis did not support G4 formation in L15.6.1 (Figure 5D) and calculated values including thermal melting temperatures (Tm), fraction folded at 37°C, and free energy changes were not significantly different regardless of monovalent cation conditions (Table S3). Neither refolding nor the presence of Mg^2+^ varied the CD spectra of L15.6.1 (Figure S10). Together, these findings support the predicted secondary structure model. However, 1D ^1^H-NMR analysis of L15.6.1 presented additional possibilities. While 1D ^1^H-NMR did not support formation of G4 structures, dispersed peaks that would support the formation of base pairing in the predicted stems were also not observed, suggesting conformational heterogeneity (Figure 5E). Thus, we cannot exclude the possibility that G4s could be a very small portion of the mixed structures, and that their frequency may have also gone undetected in our CD analysis. Of note, QGRS Mapper (*67*), a software program developed to predict the presence of G4s, predicted a potential G-quadruplex in L15.6.1 (Figure S11). Notably, the predicted structure occurs as part of the 3′ constant region, which is not essential for binding of L15.6.1 to lattice (*45*). Collectively, these results suggest that while the predicted stem-loop structures likely form, it is possible they may not be highly stable, and it is also likely that L15.6.1 is capable of assuming other structural conformations in the apo state.

Finally, as the GUGUAU motif was one of the most abundant sequence motifs identified, we sought to determine whether the motif itself was required for binding to lattice. We observed a slight but not significant decrease in binding upon mutation of the motif to CACAGA (Figure S12), suggesting that the motif may not be required for binding or that the mutations did not drastically alter aptamer structure. Notably, for five other GUGUAU-containing aptamers, the motif was also predicted to be a part of a hairpin loop or internal loop (Figure S13).

*Aptamer L15.20.1*: We next examined aptamer L15.20.1, which contains motif 3 (ACC<u>UACC</u> with a G-rich region) and was shown to bind both lattice and hexamer (Figures 2B, 3, and Table 3). None of the 3′ truncations was able to bind lattice (Figure 6A), suggesting that part or all of the 3′ constant region is required for binding. The CM generated from its cluster predicted a stem-loop structure quite different from that predicted by enzymatic and chemical probing, but that also placed the G-rich motif into a single-stranded context (Figure 6B and 6C). Mutants that disrupted the stem or the conserved YRCC motif predicted from the CM (underlined above) reduced but did not abolish binding to lattice (Figure S14), suggesting that the stem-loop may not form; however, comparisons were complicated by differences in protein-independent signal for the variants. Nuclease cleavage patterns and TMO reactivity generally matched the predicted secondary structure, but not in all positions (Figure 6C, 6D and S15).

**Figure 6:**
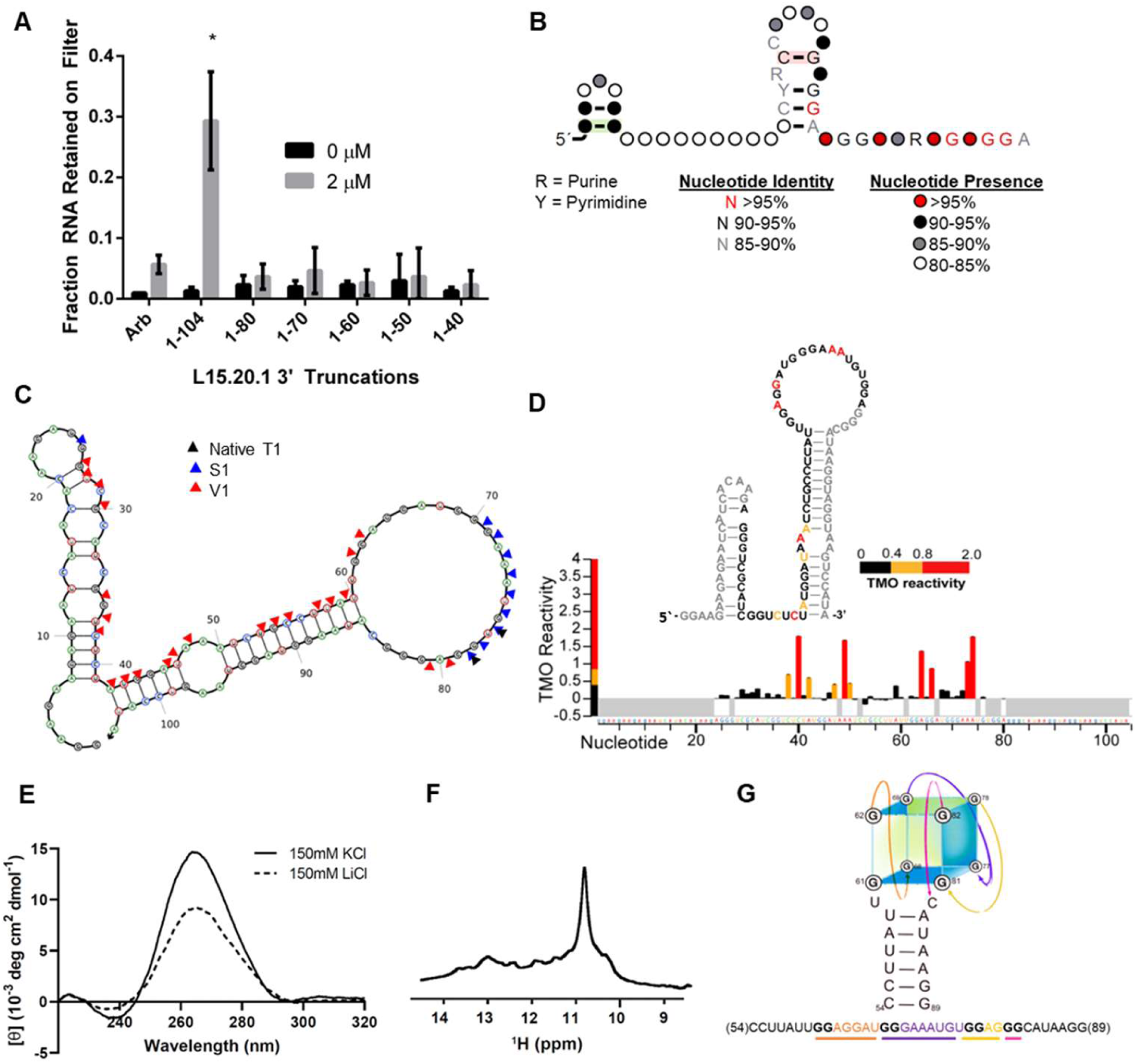
Exploration of the secondary structure of aptamer L15.20.1. (A) 3′ truncations of L15.20.1 were assessed for lattice binding using nitrocellulose filter binding assays (n = 3). Values plotted are mean ± SD.* (P < 0.05). (B) Covariance model generated from L15.20.1 and its cluster. Red shading of base means that there was no variation in the base pair. (C) Mapping the cleavage products onto the secondary structure prediction for L15.20.1 based on the covariance model. Accompanying representative gel in Figure S15. (D) Regions of TMO modification in L15.20.1 MaP experiment. Left: Predicted secondary structure of L15.20.1 where the nucleotides are colored from least (black) to most (red) TMO reactive. Nucleotides in the constant regions that were not mapped by TMO modification are colored gray. Right: The magnitude of TMO reactivity is shown at each nucleotide position. (E) CD spectra of L15.20.1 in 150 mM KCl or LiCl. (F) NMR spectrum of L15.20.1. (G) Hypothetical structure of L15.20.1 containing a G-quadruplex within the loop region, generated by artist, Miranda Tess Knutson.

Similar to L15.6.1, L15.20.1 is highly G-rich. QGRS Mapper predicted two regions within L15.20.1 that may form G4s (Figure S11). Binding to lattice was reduced by approximately 5-fold for two variants with GG→AA mutations (Figure S14) and as noted above, the 3′ constant region, which contains one of the QGRS-predicted G4s, is required for binding (Figure 6A and S11). CD spectra displayed a large increase in the G-tetrad region around 265 nm when switching from high Li^+^ to high K^+^ conditions and a minute positive region around 305 nm; both signal patterns in these two regions are associated with RNA quadruplex-like structures (Figure 6E) (*63*). UV melting at 295 nm demonstrated clear hypochromicity also generally associated with quadruplex-like structures as compared to L15.6.1 (Figure S16) (*68*). Further, a large peak was observed within the imino region of the 1D ^1^H-NMR spectrum (Figure 6F), indicating involvement of G residues in hydrogen-bonded structures. Consistent with G4 formation, Tm, fraction folded at 37°C, and calculated free energy values show evidence of increased stability in K^+^ relative to Li^+^, although this stability may be more dependent on the availability of Mg^2+^ than with the other aptamers (Table S3, Figure S17). We next examined whether lattice binding was sensitive to binding buffer conditions, as it is known that potassium ions generally stabilize G4s more than sodium ions (*69*). When we replaced the KCl with NaCl in the binding buffer, we did not observe a reduction in binding for L15.20.1 (Figure S18). This suggests either that K^+^ does not provide increased stabilization as compared to Na^+^ for L.15.20.1 or that formation of a stable G4 structure may not be required for binding. Of note, G4s with higher stability in Na^+^ as compared to K^+^ at physiological temperatures have been previously reported (*70*). While some ambiguities remain, the experimental data collectively support the possibility of G4 formation and/or non-canonical base-pairing for L15.20.1 in contrast to the CD and 1D ^1^H-NMR data for aptamer L15.6.1, which did not show any features associated with G4 formation (Figure 5E and 5F). A representation of a hypothetical G4 structure within L15.20.1 is shown in Figure 6G.

*Aptamer L15.7.1*: Aptamer L15.7.1 contains motif 1, the most common and highly G-rich motif (Figure S5). After multiple refinements of the CM in Infernal, a conserved secondary structure was not identified. However, sequence conservation of the G-rich motif among the CMs from the clusters containing this motif was high (Figure S19A-B), suggesting the motif may be important for aptamer binding to lattice. Similar to L15.20.1, 3′ truncated forms of L15.7.1 were unable to bind lattice (Figure S19C), suggesting that the 3′ region is required for binding. QGRS Mapper identified a potential G-quadruplex within L15.7.1, awarding this aptamer the highest G-score of all the aptamers (Figure S11). Notably, the predicted G-quadruplex was located in the region of the 3′ truncations and may also explain the lack of conserved secondary structure present in the CMs, as G-quadruplexes do not form traditional Watson-Crick base pairs. With this in mind, we generated a predicted secondary structure using Mfold in which the G-rich motif was forced to be single-stranded (*71*) and visualized the structure using NUPACK (*72*). Nuclease digestion patterns mapped well onto the predicted structure, with strong S1 nuclease cleavage in the region containing the G-rich motif (∼ nucleotides 51-78) (Figure S19D and S19E). CD spectra displayed an increase in ellipticity at ∼265 nm under high K^+^ conditions, in contrast to that of Li^+^ (Figure S19F), further supporting G-quadruplex formation. The 1D ^1^H-NMR imino proton spectra suggested that G-tetrad motifs are possible but may be temperature dependent, as signal was observed only at 35°C (Figure S19G). Calculated values reveal higher folding and greater stability under high K^+^ conditions (Table S3, Figure S20). When we replaced the KCl with NaCl in our binding assay or mutated G to A within L15.7.1, we observed reduced lattice binding, further supporting formation of G4 structures and suggesting the structure may be important for lattice binding (Figure S18 and S21). Collectively, these data support preliminary assignment of a G-quadruplex structure to this region.

*Aptamer H7.10.1*: Considering that aptamer H7.10.1 binds both lattice and hexamer and most aptamers in the libraries cannot bind hexamer, it was not surprising that H7.10.1 showed no sequence relationship to the motifs found in the top 1000 cluster seed sequences from L15 or the top 1000 cluster seed sequences from L17+H7. A truncation containing nucleotides 1-80 retained lattice binding to the same degree as the full-length aptamer, while binding was slightly reduced upon further truncation to 1-70 and 1-60 (Figure S22A). When a CM generated from the alignment of the 51 sequences from the H7.10.1 cluster (Figure S22B) was searched against the top 1000 cluster seed sequences from L15 or L17+H7 using Infernal, no other cluster seed sequences matched. Nuclease probing (Figure S22C) mapped well onto the predicted stem-loop structure (Figure S22D). However, QGRS Mapper predictions, as well as CD and 1D ^1^H-NMR data may support the potential formation of G4s or non-canonical base pairing. The CD spectra displayed an ellipticity increase in the G-tetrad region under high K^+^ conditions when compared to that of Li^+^ (Figure S22E), and the 1D ^1^H-NMR imino proton spectrum also supported the possibility of G4 formation or non-canonical base pairing (Figure S22F). Further, calculated values for thermal stability, fraction folded, and free energy improved under high K^+^ conditions relative to Li^+^, and overall Tm in K^+^ is higher for this aptamer than for any of the others (Table S3). Interestingly, under high K^+^ conditions, there appear to be structural differences between the aptamer’s refolded and native states (Figure S23). Collectively, these data support preliminary assignment of a G-quadruplex structure.

### Amenability of aptamers for affinity purification

To facilitate development of these aptamers as tools to study CA assembly forms in cells, we next evaluated whether our candidate aptamers could be adapted for affinity purification of CA from cell lysates. Aptamers L15.6.1 and H7.10.1 retained binding to lattice in the absence of their 3′ constant regions (Figure S22A and (*45*)), suggesting the 3′ end may be amenable to modification for affinity purification. When we annealed an oligonucleotide complementary to the 3′ constant region, we found that oligo annealing abolished binding for all aptamers except L15.6.1 (Figure 7A). Based on these results, we hypothesized that when annealed to a biotinylated oligo on the 3′ end, L15.6.1 would pull down lattice from cell lysates, while H7.10.1 would not. Indeed, we observed pulldown of lattice from cell lysates for L15.6.1, but not H7.10.1 (Figure 7B and 7C). Notably, L15.6.1 was unable to pull down hexamer (data not shown). Thus, while some aptamers will require additional structural investigation and optimization for use as affinity purification reagents, some aptamers may be easily modified for this application.

**Figure 7.**
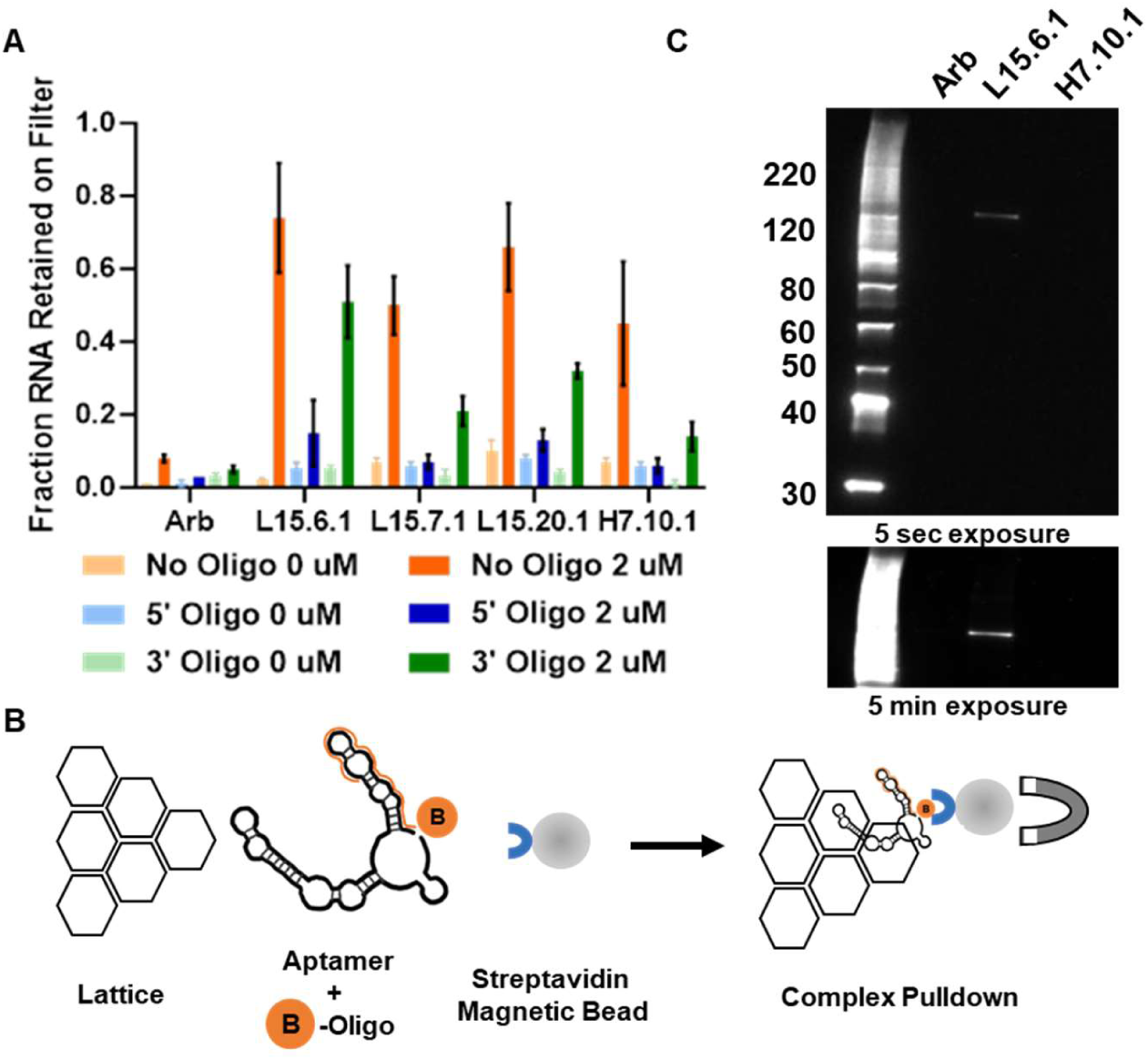
Aptamer-mediated recovery of CA from cell lysates. (A) Binding to lattice (n = 3) was accessed for aptamers annealed to an oligonucleotide complementary to the 3′ constant region using nitrocellulose filter binding assays. Values plotted are the mean ± SD. (B) Schematic of pulldown protocol (not drawn to scale). (C) The ability of aptamers annealed to a biotinylated oligonucleotide complementary to the 3′ constant region to pulldown lattice spiked into cellular lysates was determined using streptavidin magnetic bead separation followed by western blot detection of CA using anti-p24 antibody (n=2).

## DISCUSSION

The differentiation selection strategy described here demonstrates the utility of combining aptamer selections with HTS and subsequent bioinformatic analyses to identify aptamer subsets with specialized binding phenotypes from within an already highly converged library, L15. Approximately 67% of the HTS reads in L15 were the top twenty most abundant sequences. Most of these top sequences were not predicted to be hexamer binders due to their strong depletion from L15 to L17+H7, and all aptamers that were predicted to be lattice-specific based on their trajectory-specific enrichment patterns were indeed determined experimentally to be lattice-specific binders. For a selection in which the aptamer pool is exposed to multiple potential targets, strong depletion against one or more specific targets appears to be an acceptable indicator that the aptamer likely does not bind the target. In contrast, binding predictions were less successful for aptamers that had enrichment values near 1 (i.e., fractional representation in the population did not change) or greater in the lattice and hexamer trajectory and were, therefore, predicted to be capable of binding both lattice and hexamer. This is likely because there were not many hexamer-binding sequences present in the highly converged L15 library used as the branch point. As it was likely that L15 contained a low abundance of hexamer binders due to convergence, the fact that aptamers capable of binding both lattice and hexamer were identified demonstrates the power of the differentiation selection strategy used here. Indeed, our results suggest that binders in low abundance can be parsed out from highly converged aptamer populations if they are present. Adaptations of this strategy could be applied to identify additional CA assembly state binders, as well as aptamers capable of differentiating among other protein targets.

A higher number of hexamer binders may have been present in pre-enriched but less converged rounds of the initial selection (e.g. L8). Aptamer libraries typically converge after five to fifteen rounds of selection (*73*). HTS analysis of several selection rounds would be beneficial in identifying branch points for differentiation selections. Ideally, the branch point for pursuing specialized recognition would occur at a round where some aptamer enrichment was observed but had not completely converged. Notably, it is difficult to suggest specific parameters since sequence distributions are dependent on the selection and will vary across aptamer libraries and targets. With respect to our selection, L15 had seven sequences with RPM values greater than 50,000 that made up nearly 50% of the total reads from the library. In contrast, L8 had one sequence with an RPM value greater than 50,000, and the ten most abundant sequences from L8 made up almost 30% of the total reads. Whether L8 or earlier rounds may contain additional hexamer binders not identified from L15 remains to be determined.

Incorporation of a hexamer depletion trajectory was not informative in predicting binding phenotypes, likely due to the high convergence of L15. In contrast, incorporation of an enrichment selection trajectory using lattice and hexamer and a nitrocellulose depletion trajectory improved our ability to interpret the HTS data from the differentiation selection. As every RNA sequence from L15 had the opportunity to pass through or bind to the nitrocellulose filter, enrichment values from these trajectories could be compared to identify false positive hits. While most sequences did not show substantial enrichment in the nitrocellulose trajectory, L15.45.1 had enriched approximately 20-fold in the nitrocellulose trajectory and 66-fold in the positive hexamer trajectory. Because of its enrichment within the nitrocellulose trajectory, L15.45.1 was not predicted to bind hexamer and this was confirmed experimentally. To improve the utility of the nitrocellulose depletion trajectory, it may be worthwhile to increase the number of depletion rounds corresponding to the number of enrichment rounds. Then, if sequences did not bind hexamer but did bind nitrocellulose, the enrichment values for these sequences should be comparable.

Differentiation selection identified four candidates for structural analysis. Biochemical investigations suggested that CA-binding aptamers may contain different structural features and that each of the aptamers may adopt an ensemble of structures independent of protein binding. L15.6.1 likely exhibits substantial conformational heterogeneity while L15.20.1 appears to contain a G4 structure. Results for aptamers L15.7.1 and H7.10.1 were less conclusive but were also suggestive of G- or mixed-quadruplexes or non-canonical base pairing. Some of the potential structures supported by our data may be important for binding to lattice, while others may form, but not impact lattice binding. Notably, aptamer structures displayed differential sensitivity to experimental conditions, including temperature, highlighting the importance of selecting aptamers using conditions appropriate to the desired application. Future studies will focus on better understanding apo and protein-bound aptamer structures, including in cells, and identifying aptamer binding sites on CA.

Notably, all four aptamers tested were able to bind stabilized mature capsid cores, suggesting that these aptamers may be capable of interacting with a biologically relevant form of CA in cells in addition to the *in vitro* assembled lattice used as a selection target. While some aptamers that interact with CA assembly forms in cells could alter virus replication, others may have no biological impact. Both outcomes would be beneficial to the study of CA assembly forms. Current evidence suggests that some CA-binding aptamers affect virus infectivity while others do not [manuscript in preparation and (*45*)]. We previously observed that L15.6.1 (CA15-2) inhibited infection when expressed in virus producing cells but was unable to protect target cells from new infection (*45*). Together with the ability of L15.6.1 to bind the mature capsid core, these collective results support development of L15.6.1 (and other aptamers) as an affinity purification tool for identification of CA-interacting host factors in target cells or to track mature cores in target cells using microscopy. Indeed, L15.6.1 was easily modified for pulldown of lattice spiked into cell lysates using a complementary biotinylated oligo annealed to its 3′ constant region. Other aptamers were not as easily modified, as oligo annealing abolished their binding to CA, and will thus require further optimization. Aptamers that impact infectivity could be useful to perturb CA-host factor interactions, identify biologically important sites on CA for therapeutic targeting, or to study the functional contributions of different CA assembly forms to replication processes in cells.

In conclusion, we have successfully identified aptamers with distinct binding specificities for HIV-1 capsid assembly forms. While most of the identified aptamers specifically bind lattice, we were also able to identify aptamers capable of binding both lattice and hexamer from our highly enriched L15 library. This study supports the feasibility of identifying aptamers with specificity for different higher order protein assemblies and provides new tools for the study of capsid assembly forms in cells. Furthermore, our study highlights the power of combining a differentiation selection approach with HTS analysis to identify low abundance sequences from highly converged libraries. Finally, and importantly, our structural studies highlight challenges in determination of apo aptamer RNA structure, the importance of experimental conditions in structure determination and aptamer development, and the utility of combining a variety of different techniques.

## DATA AVAILABILITY

All data are available in the main text or the supplementary materials.

## SUPPLEMENTARY DATA

Supplementary Data are available at NAR online.

## AUTHOR CONTRIBUTIONS

Paige Gruenke: Conceptualization, Formal analysis, Methodology, Validation, Visualization, Writing—original draft. Miles Mayer: Formal analysis, Visualization, Writing—review & editing. Rachna Aneja: Formal analysis, Methodology, Writing—review & editing. Zhenwei Song: Formal analysis, Methodology, Writing—review & editing. Xiao Heng: Formal analysis, Methodology, Writing—review & editing. Donald Burke: Methodology, Writing—review & editing. Margaret Lange: Conceptualization, Formal analysis, Methodology, Visualization, Writing—review & editing.

## ACKNOWLEDGEMENTS

The authors would like to acknowledge tremendously helpful contributions by Drs. Stefan Sarafianos, Owen Pornillos, and Barbie Ganser-Pornillos, whose labs supplied constructs for protein expression, purified protein, aided in establishing protein purification methodologies in the Lange laboratory, and were always available to discuss results or provide helpful suggestions. We also acknowledge Miranda Tess Knutson, who provided an artistic rendering for Figure 6G.

