## Supplemental Data for "Differentiation SELEX approach identifies RNA aptamers with different specificities for HIV-1 capsid assembly forms"

Paige R. Gruenke *et al*.

**This PDF file includes:**

Figs. S1 to S#23
Tables S1 to S#3

**Fig. S1**

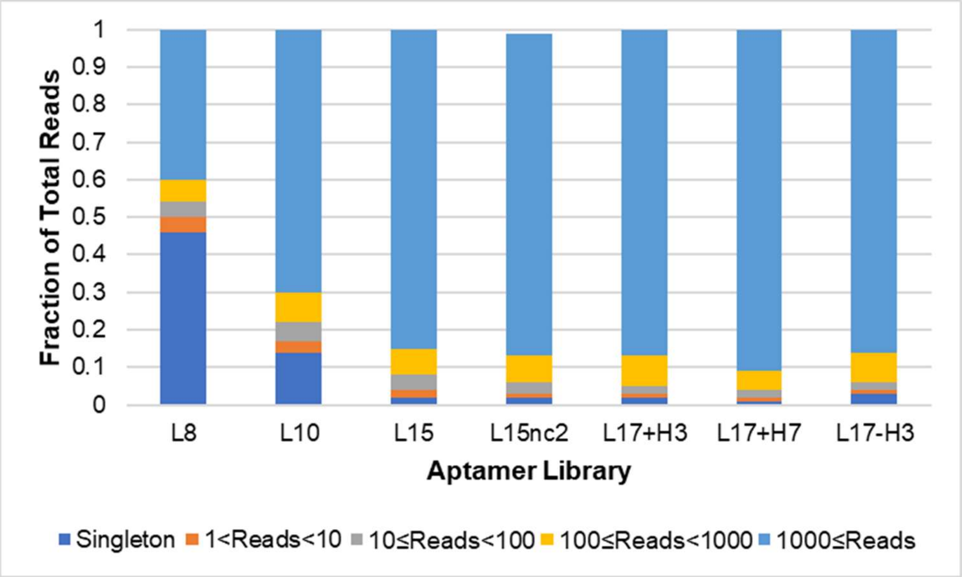


**Figure S1: Binned sequence abundance plot for CA aptamer libraries.** Unique sequences were binned according to their read counts for each aptamer library, and the fraction of total reads for each aptamer bin was plotted.

**Fig. S2.**

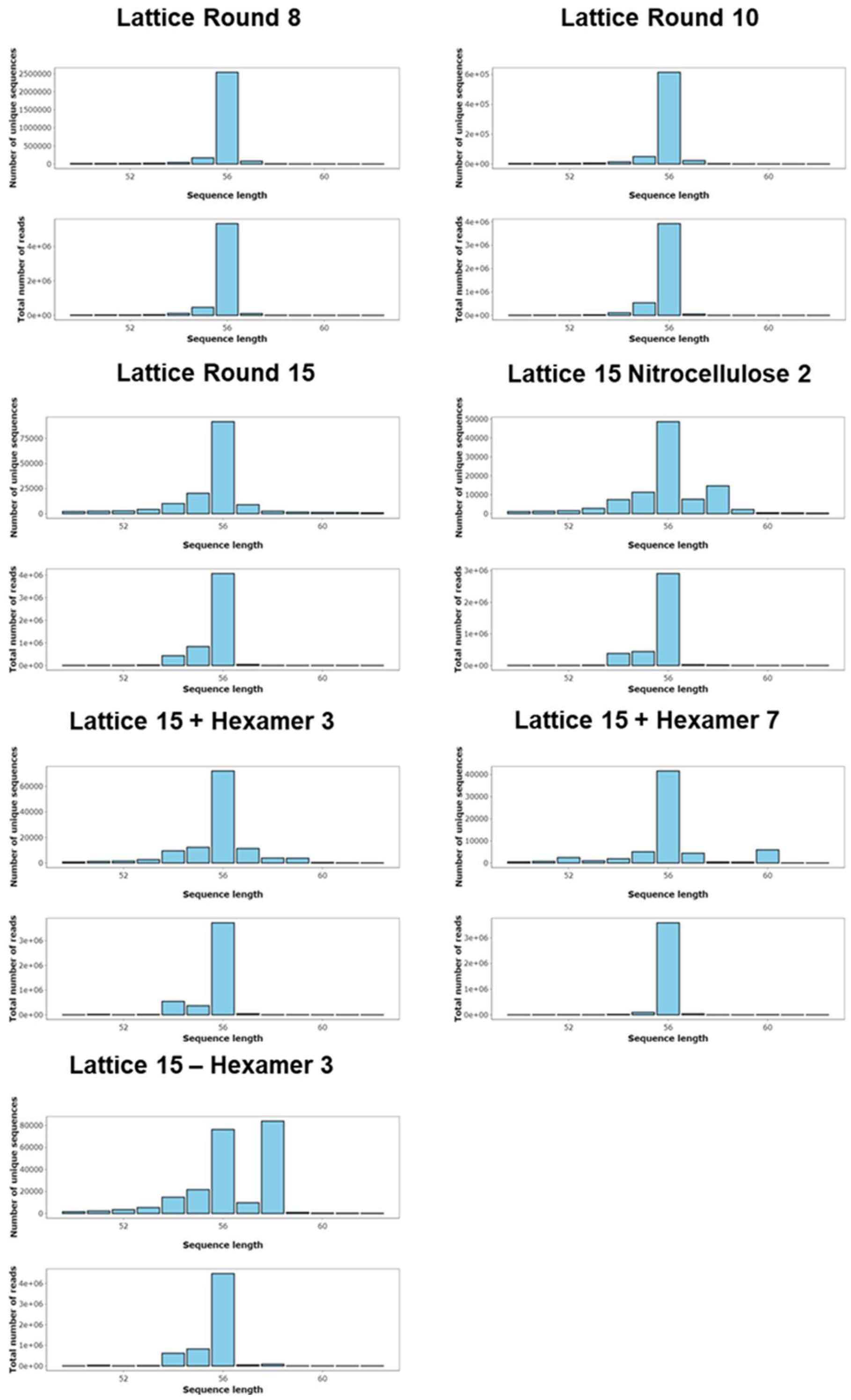


**Figure S2: Sequence length histograms of the CA aptamer library.** For each library specified above, the number of unique sequences and the total number of reads were plotted at each sequence length.

Fig. S3.

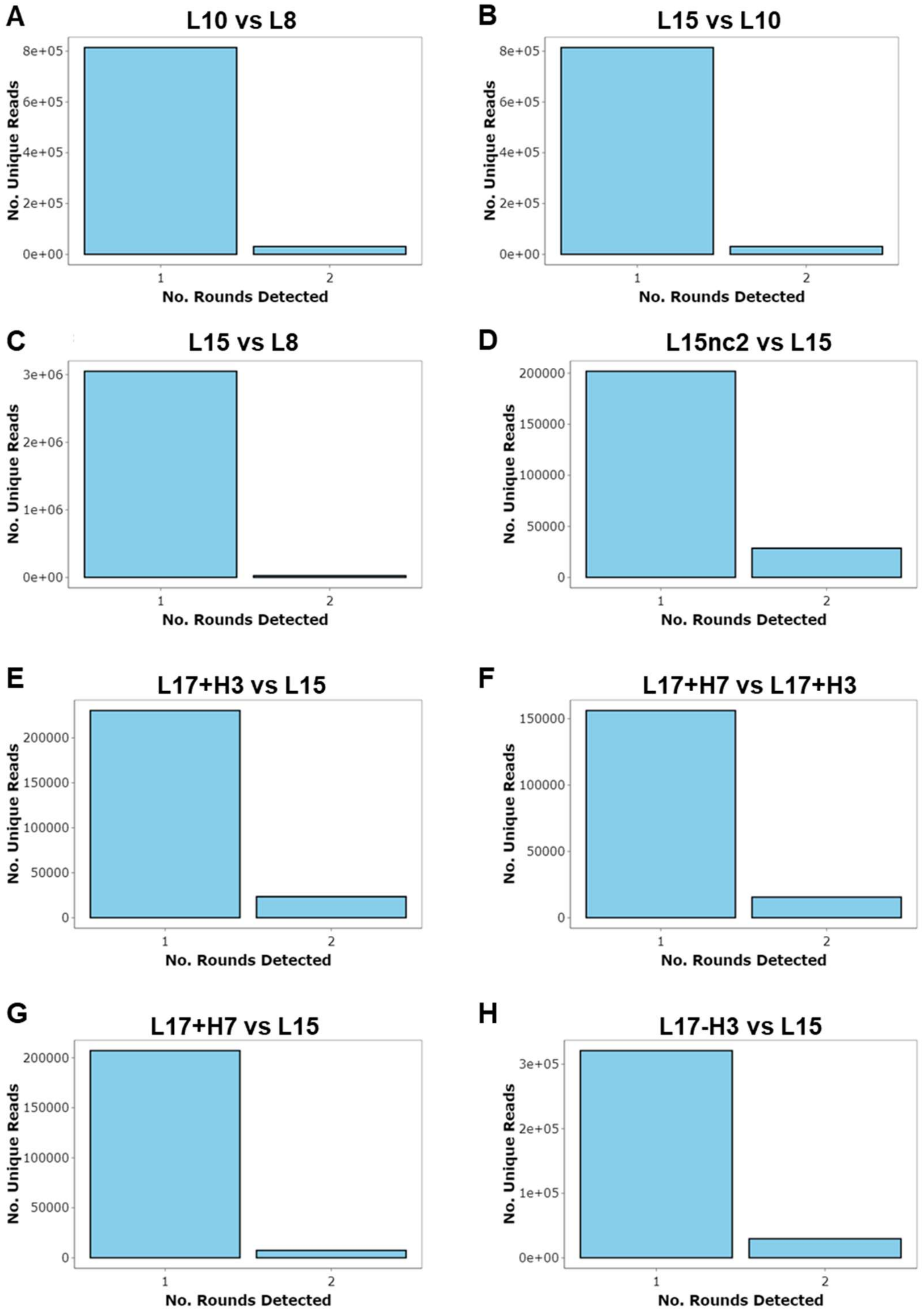


**Figure S3: Sequence persistence between CA aptamer libraries.** The number of unique sequences present in either one or both aptamer libraries were plotted.

Fig. S4.

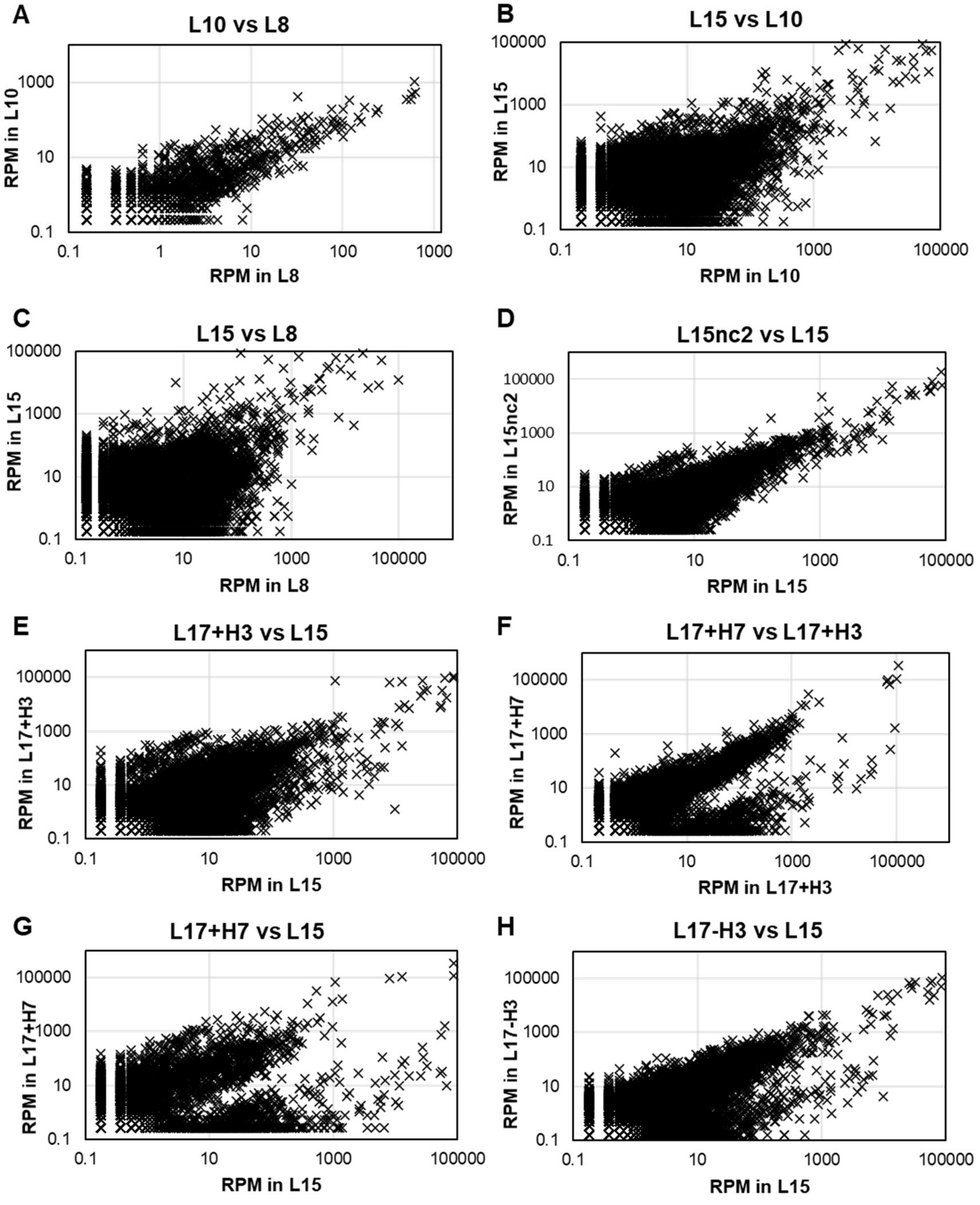


**Figure S4: Comparison of sequence read frequencies among CA aptamer libraries.** Sequences were plotted based on their read frequencies in both aptamer libraries in reads per million (RPM). Only sequences that were present in both libraries were plotted. Both axes are plotted on a logarithmic scale.

**Fig. S5**

**Motif 1**

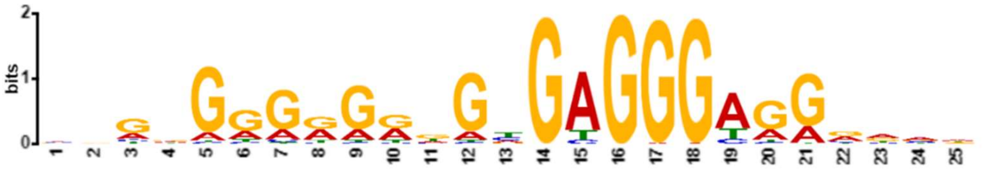


**Motif 2**

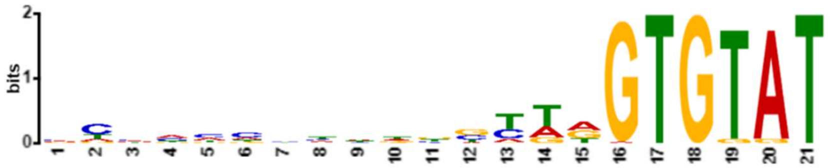


**Motif 3**

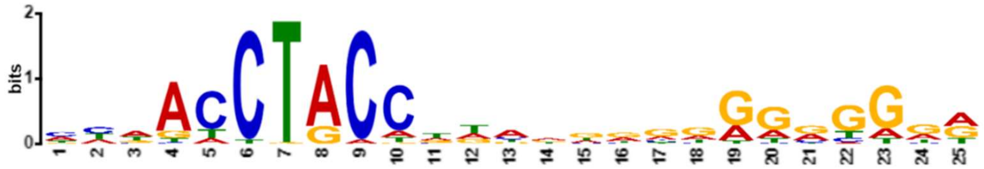


**Figure S5: Sequence logos for the top 3 most abundant sequence motifs identified using MEME Suite.**

**Fig. S6**

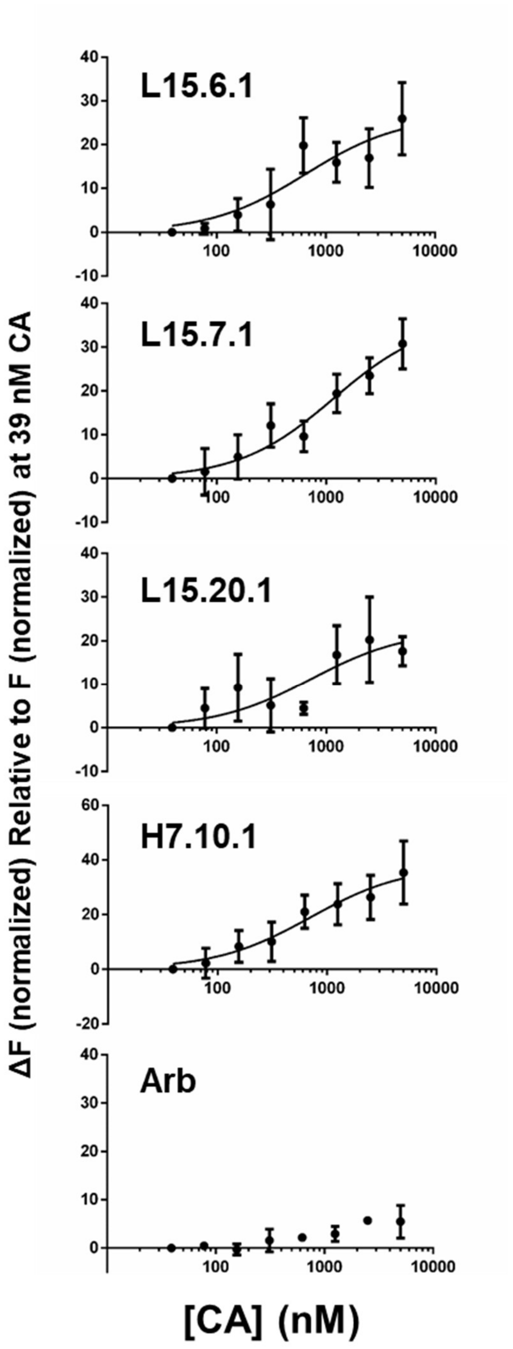


**Figure S6: Apparent Kd Binding Curves for Representative Aptamers to CA Lattice.** Binding curves from microscale thermophoresis (n = 4). RNAs were generated with a 3′ tail extension and annealed to a Cy5-labeled anti-tail oligonucleotide in the presence of increasing concentrations of protein. For binding assays, protein concentration is based on the CA monomer.

**Fig. S7**

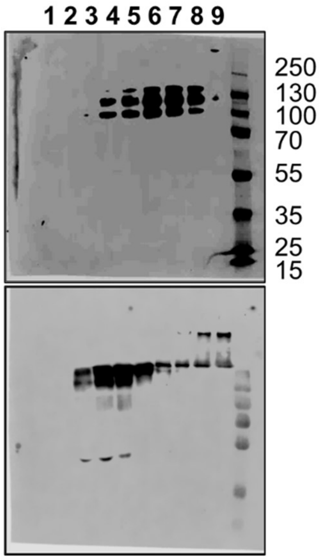


**Figure S7: Purification of A14C/E45C stabilized viral cores.** The pellet (lane 9) contained the stabilized capsid cores.

**Fig. S8**

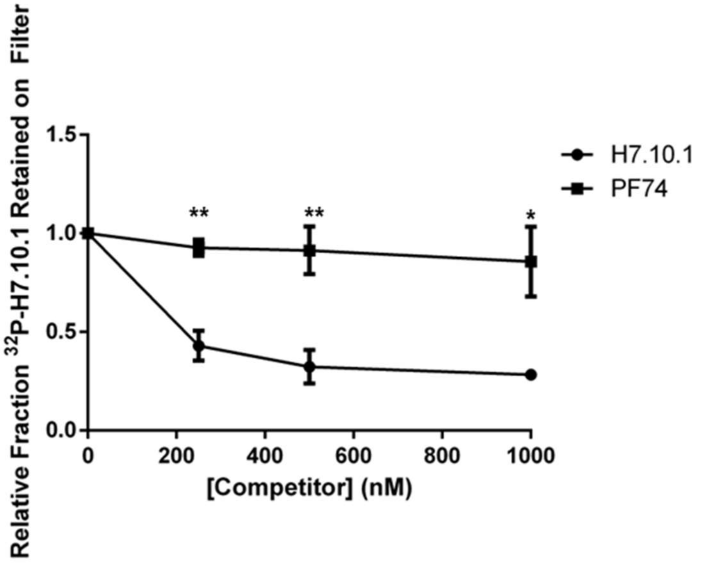


**Figure S8: Competition of unlabeled aptamer H7.10.1 or PF74 and radiolabeled aptamer H7.10.1 for binding to CA lattice.** The relative fraction of radiolabeled aptamer H7.10.1 retained on the nitrocellulose filter was determined using full length unlabeled H7.10.1 and PF74 as competitors ($n = 3$). Values are the mean ± SD. * ($P < 0.05$), ** ($P < 0.01$).

**Fig. S9**

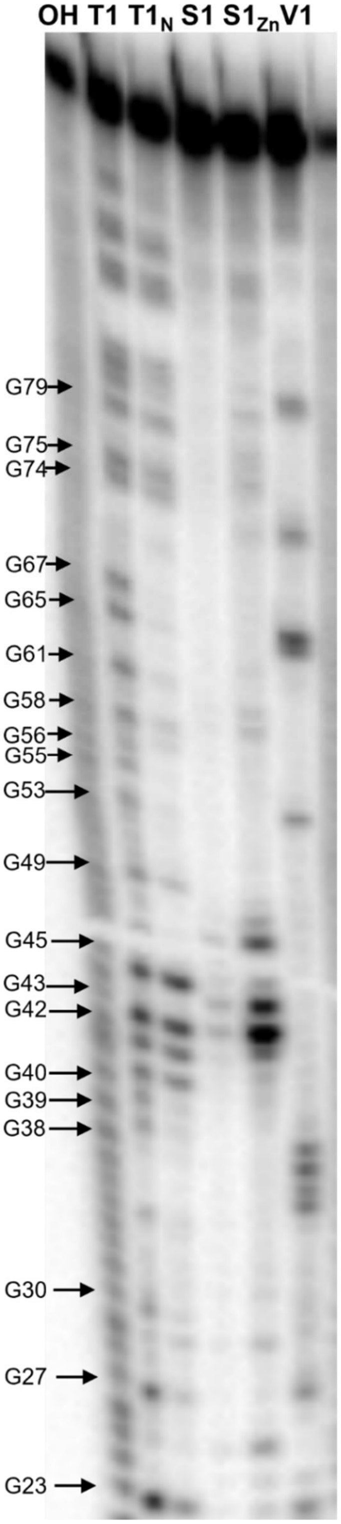


**Figure S9. Nuclease digestion profiling for L15.6.1.**

**Fig. S10**

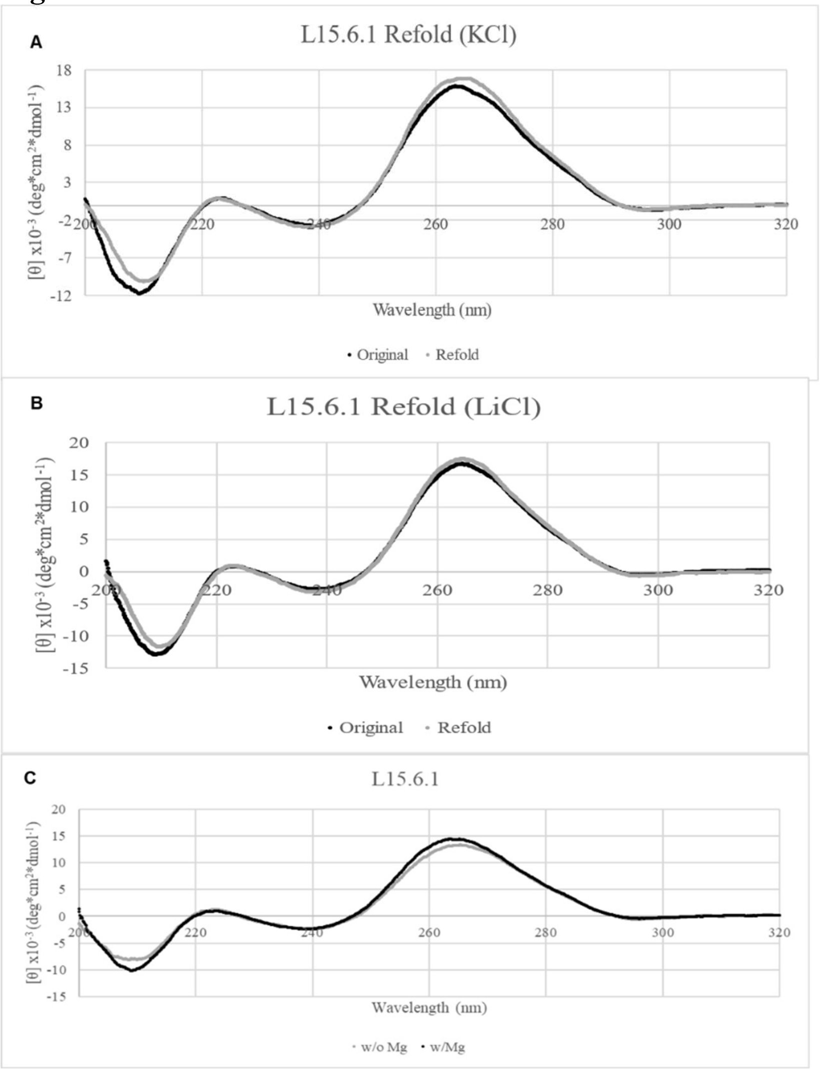


**Figure S10: L15.6.1 Refolding and Magnesium Dependence.** (A) Refolding in KCl. (B) Refolding in LiCl. (C) Magnesium Dependence of L15.6.1.

**Fig. S11**

| Aptamer | Position | Length | QGRS | G-Score |
|---|---|---|---|---|
| L15.6.1 | 74 | 21 | GGCAUGCGGGCAUAAGGUAGG | 17 |
| L15.7.1 | 53 | 30 | GGGAAAGGAGUGGGUGGUACGGGACAUGGG | 38 |
| L15.20.1 | 44 | 27 | GGAUAAAUCUGCCUUAUUGGAGGAUGG | 6 |
| | 78 | 17 | GGAGGGCAUAAGGUAGG | 18 |
| H7.10.1 | 26 | 27 | GGCGGGUAAACUAAGUCCUUGGGAGGG | 9 |
| | 75 | 20 | GGAUGGGGGCAUAAGGUAGG | 18 |
| L15.1.1 | NA | NA | None Identified | 0 |
| Arbitrary | NA | NA | None Identified | 0 |

```
L15.6.1     GGAAGAAGAG AAUCAUACAC AAGAUCGACG UACCUCAGGG UGGUGUAUGA CUGAGGUGAA
GACUGUGAAC CAUGGCAUGC GGGCAUAAGG UAGGUAAGUC CAUA

L15.7.1     GGAAGAAGAG AAUCAUACAC AAGACCAACC AGAGACGCAC CAGAGUCCUC GAGGGAAAGG
AGUGGGUGGU ACGGGACAUG GGGCAUAAGG UAGGUAAGUC CAUA

L15.20.1    GGAAGAAGAG AAUCAUACAC AAGAGGGUCG CAUCGGUCUC UAUGGAUAAA UCUGCCUUAU
UGGAGGAUGG GAAAUGUGGA GGGCAUAAGG UAGGUAAGUC CAUA

H7.10.1     GGAAGAAGAG AAUCAUACAC AAGAUGGCGG GUAAACUAAG UCCUUGGGAG GGAGCGAUGG
UUCUUAGCCU CGAGGGAUGG GGGCAUAAGG UAGGUAAGUC CAUA

L15.1.1     GGAAGAAGAG AAUCAUACAC AAGAGCCGAU GCAGUCCCAA UGUCGUAUGA GAUGUGACUA
CCUACCACUG UUUAGUGUAU GGGCAUAAGG UAGGUAAGUC CAUA

Arbitrary   GGGAAAAGCG AAUCAUACAC AAGAAAUUUG GACUUUCCGC CCUUCUUGGC CUUUAUGAGG
AUCUCUCUGA UUUUUCUUGC GUCGAGUUUU CCGGGGGCAU AAGGUAUUUA AUUCCAUA
```

**Figure S11: Prediction of potential G-quadruplex structures within aptamer sequences using QGRS Mapper.** Aptamer constant regions are highlighted in yellow and predicted G-quadruplexes are bolded and underlined.

**Fig. S12**

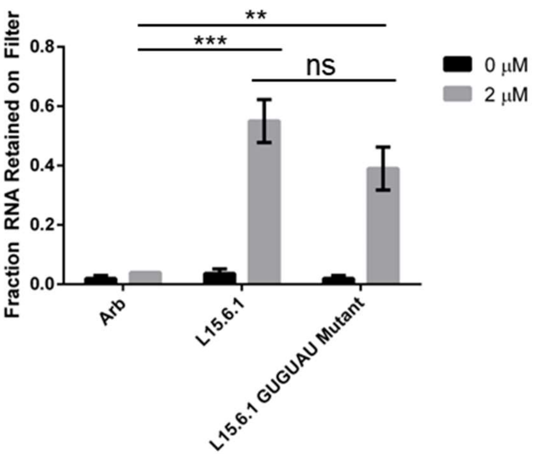


**Figure S12: Binding of L15.6.1 GUGUAU mutant to CA lattice.** The GUGUAU motif in L15.6.1 was mutated to CACAGA, and binding to CA lattice was assessed using nitrocellulose filter binding assays (n = 3). Values are the mean ± SD. ns ($P > 0.05$), ** ($P < 0.01$), *** ($P < 0.001$).

**Fig. S13**

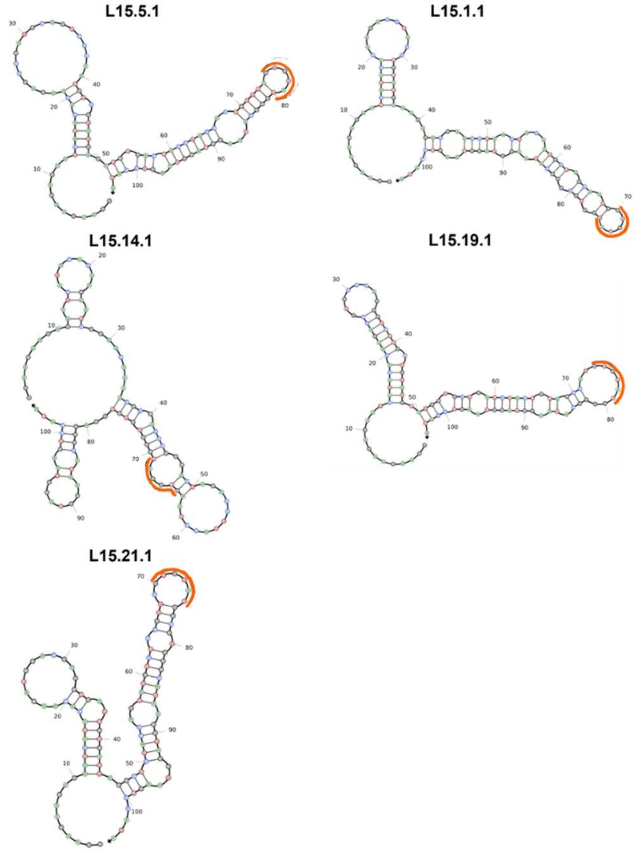


**Figure S13: Predicted secondary structures of other GUGUAU-containing aptamers.** For the aptamers shown above, the predicted secondary structures generated using NUPACK (72) are shown, and the GUGUAU motif is marked with an orange line.

**Fig. S14**

**A**

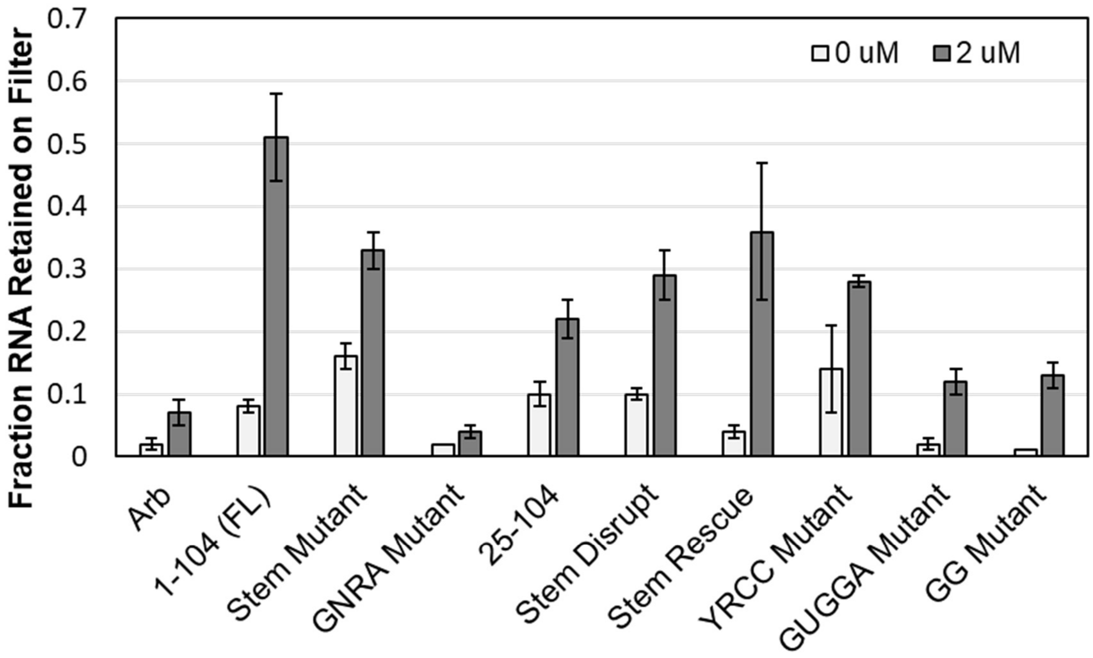


**B**

**L15.20.1**

GGGUCGCAUCGGUCUCUAUGGAUAAAUCUGCCUUAUUGGAGGAUGGGAAAUGUGGA

**L15.20.1 Stem Mutant**

GGGUCGCAUCGGUCUCUAUGGAU**UU**AUCUGCCUUAUUGGAGGAUGGGAAAUGUGGA

**L15.20.1 GNRA Mutant**

GGGUCGCAUCGGUCUCUAUGGAUAAAUCUGCCUUAU**GUAA** **+ remove nucleotides 81-84**

**L15.20.1 Stem Disrupt**

GGGUCGCAUCGGUCUCUAUGGAUAAA**AAC**GCCUUAUUGGAGGAUGGGAAAUGUGGA

**L15.20.1 Stem Rescue**

GGGUCGCAUCGGUCUCUAUGGAUAAA**AAC**GCCUUAUUGG**GUU**AUGGGAAAUGUGGA

**L15.20.1 YRCC Mutant**

GGGUCGCAUCGGUCUCUAUGGAUAAAUCU**UAA**UUAUUGGAGGAUGGGAAAUGUGGA

**L15.20.1 GUGGA Mutant**

GGGUCGCAUCGGUCUCUAUGGAUAAAUCUGCCUUAUUGGAGGAUGGGAAAU**A**U**AA**A

**L15.20.1 GG Mutant**

GGGUCGCAUCGGUCUCUAUGGAUAAAUCUGCCUUAUUGGAGGAU**AA**GAAAUGUGGA

**Figure S14: Assessing Binding of L15.20.1 Variants for Binding to CA Lattice.** (A) Binding was assessed using nitrocellulose filter binding assays. (n = 3) (B) Mutations that were introduced into L15.20.1 are shown bolded and with yellow highlight.

**Fig. S15**

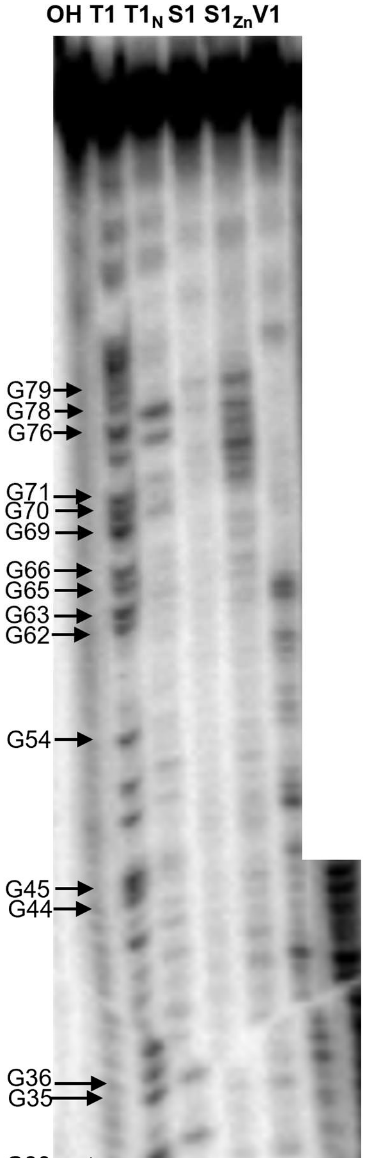


**Figure S15. Nuclease digestion profiling for L15.20.1.**

**Fig. S16**

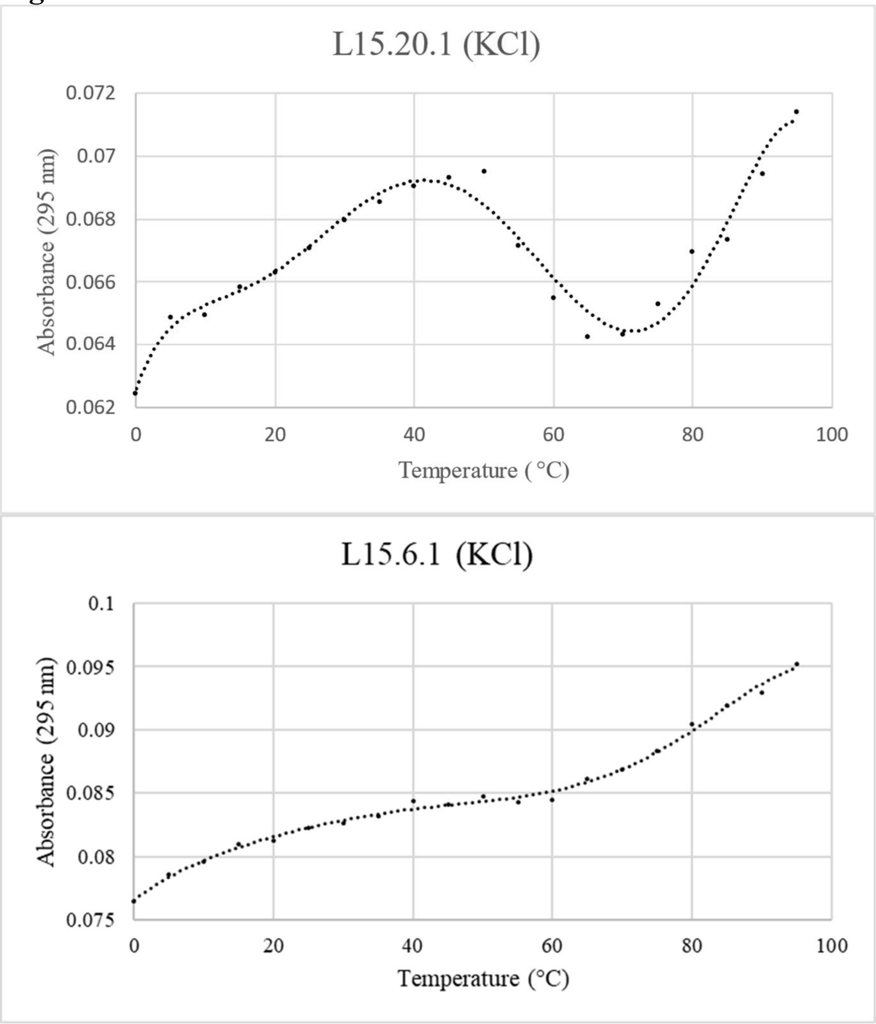


**Figure S16: Hypochromicity.** L15.20.1, predicted to form a G-quadruplex structure, displays clear hypotchromicity as compared to L15.6.1, which was not predicted to form a G-quadruplex structure.

**Fig. S17**

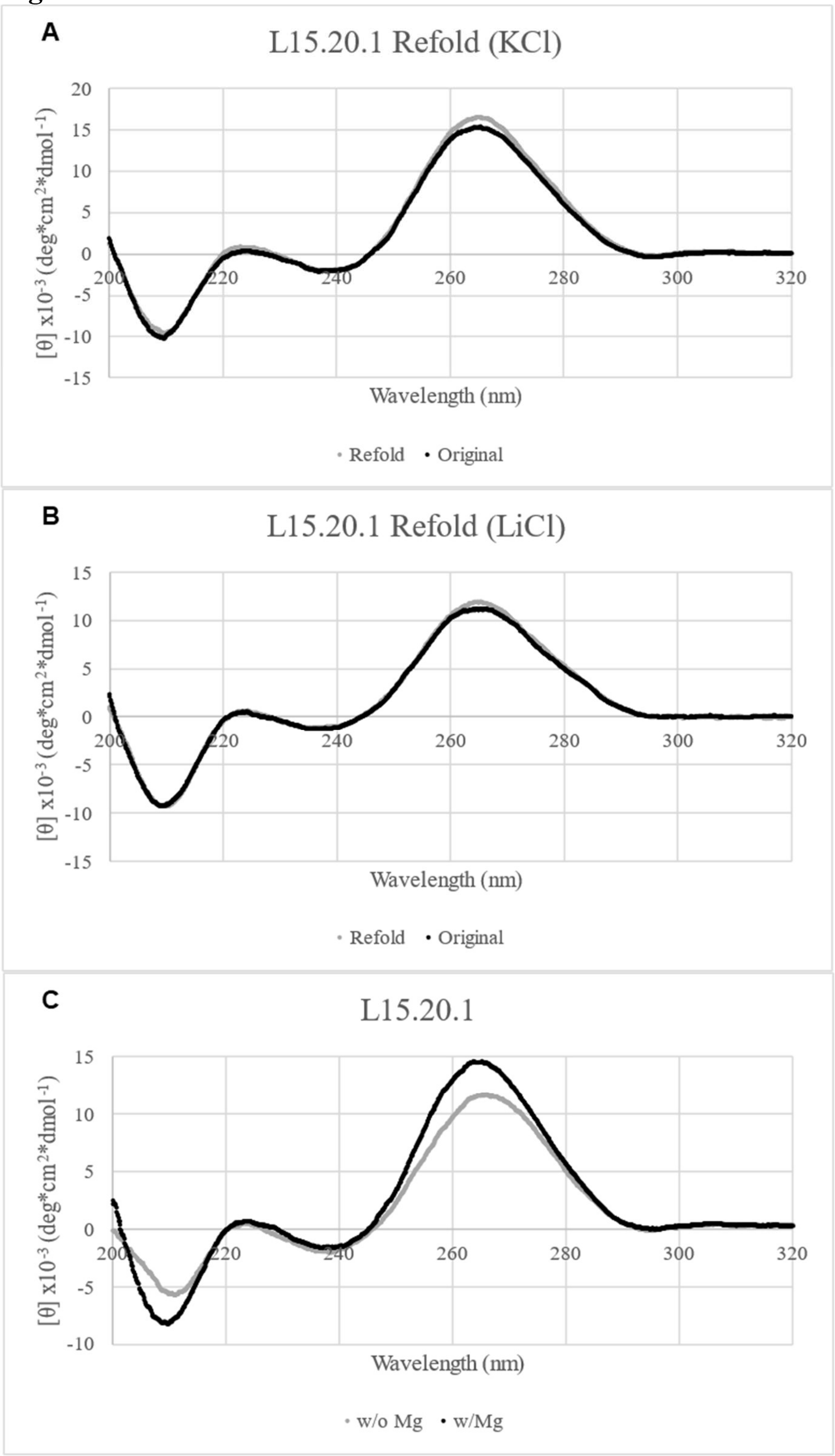


**Figure S17: L15.20.1 Refolding and Magnesium Dependence.** (A) Refolding in KCl. (B) Refolding in LiCl. (C) Magnesium Dependence of L15.20.1.

**Fig. S18**

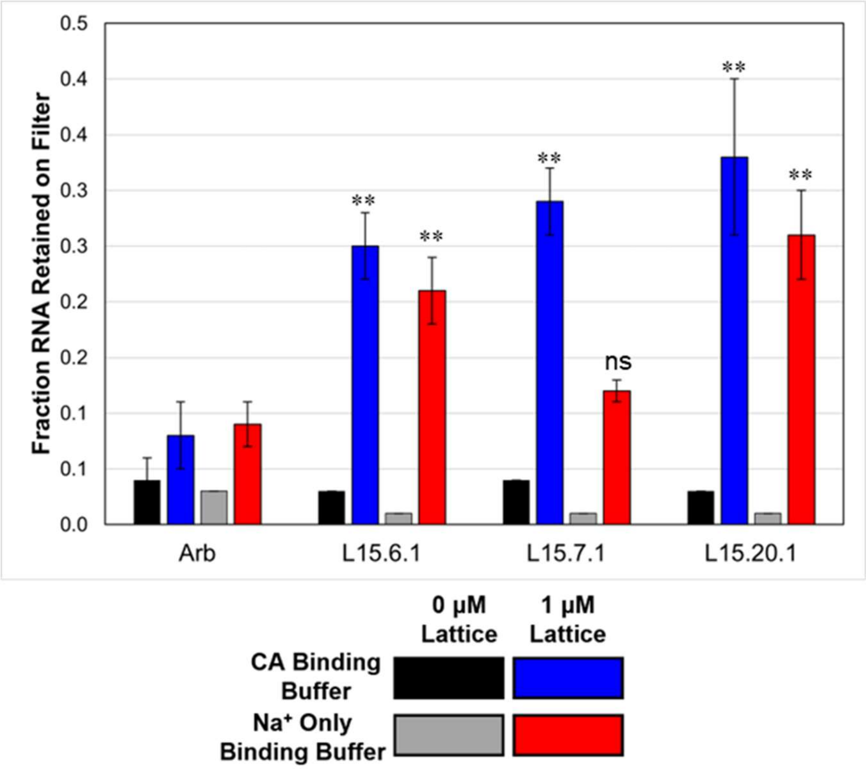


**Figure S18: Assessing binding of aptamers to CA lattice in different binding buffers.** Nitrocellulose filter binding assays were performed to determine whether aptamer binding was affected when the potassium ions in the capsid binding buffer were replaced with sodium ions (n = 3). Values are mean ± SD. Statistical comparisons were made against the binding values of the arbitrary control in the same buffer. ** (P<0.01), ns (P>0.05).

Fig. S19

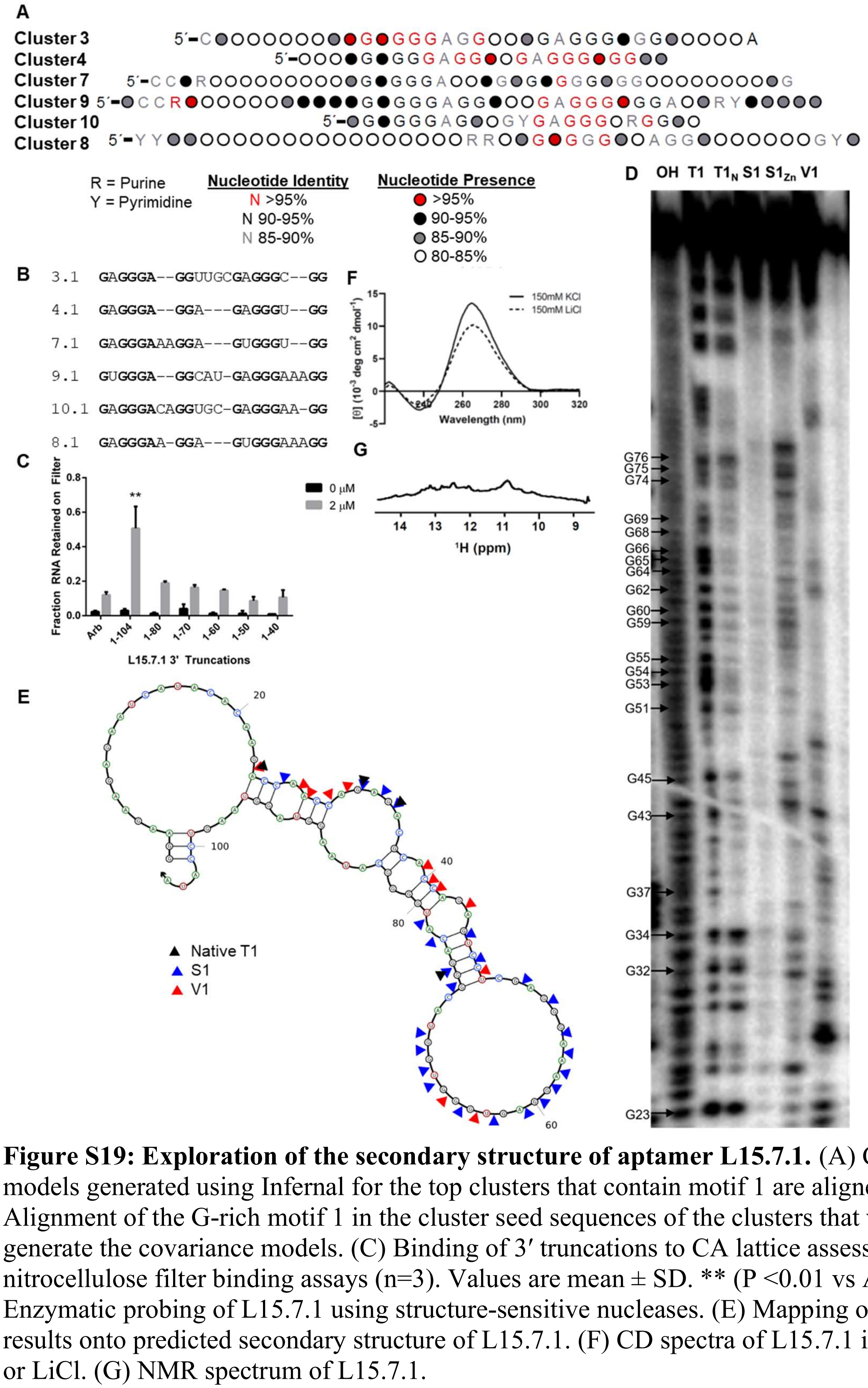


**Figure S19: Exploration of the secondary structure of aptamer L15.7.1.** (A) Covariance models generated using Infernal for the top clusters that contain motif 1 are aligned. (B) Alignment of the G-rich motif 1 in the cluster seed sequences of the clusters that were used to generate the covariance models. (C) Binding of 3′ truncations to CA lattice assessed using nitrocellulose filter binding assays (n=3). Values are mean ± SD. ** (P <0.01 vs Arb). (D) Enzymatic probing of L15.7.1 using structure-sensitive nucleases. (E) Mapping of digestion results onto predicted secondary structure of L15.7.1. (F) CD spectra of L15.7.1 in 150 mM KCl or LiCl. (G) NMR spectrum of L15.7.1.

**Fig. S20**

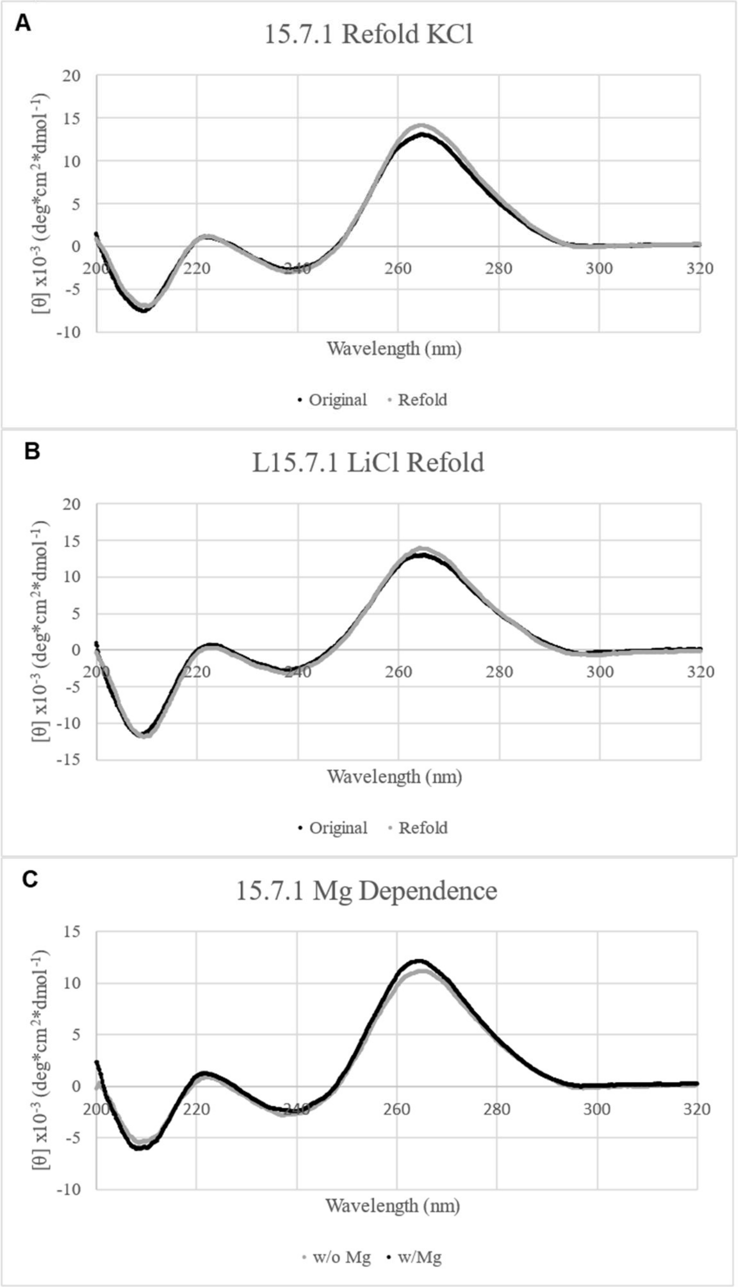


**Figure S20: L15.7.1 Refolding and Magnesium Dependence.** (A) Refolding in KCl. (B) Refolding in LiCl. (C) Magnesium Dependence of L15.7.1.

**Fig. S21**

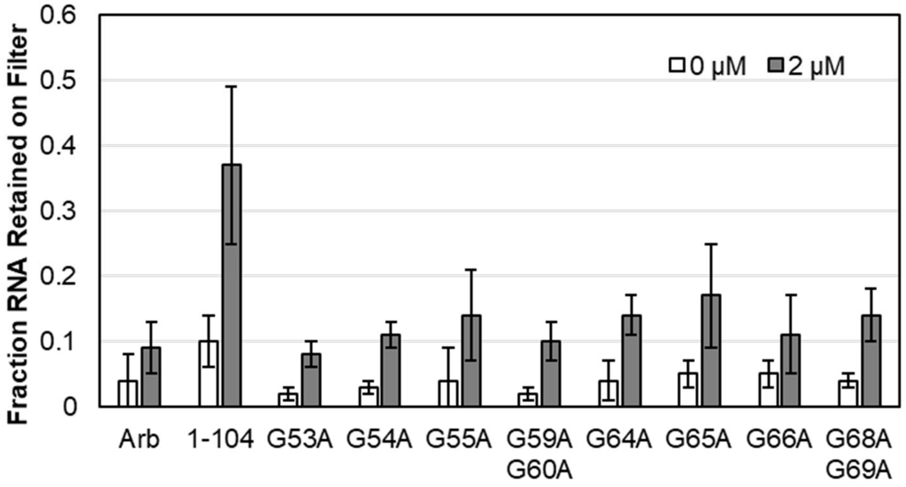


**Figure S21: Assessing Binding of L15.7.1 G to A Mutants for Binding to CA Lattice.** (A) Binding was assessed using nitrocellulose filter binding assays ($n \geq 3$).

**Fig. S22**

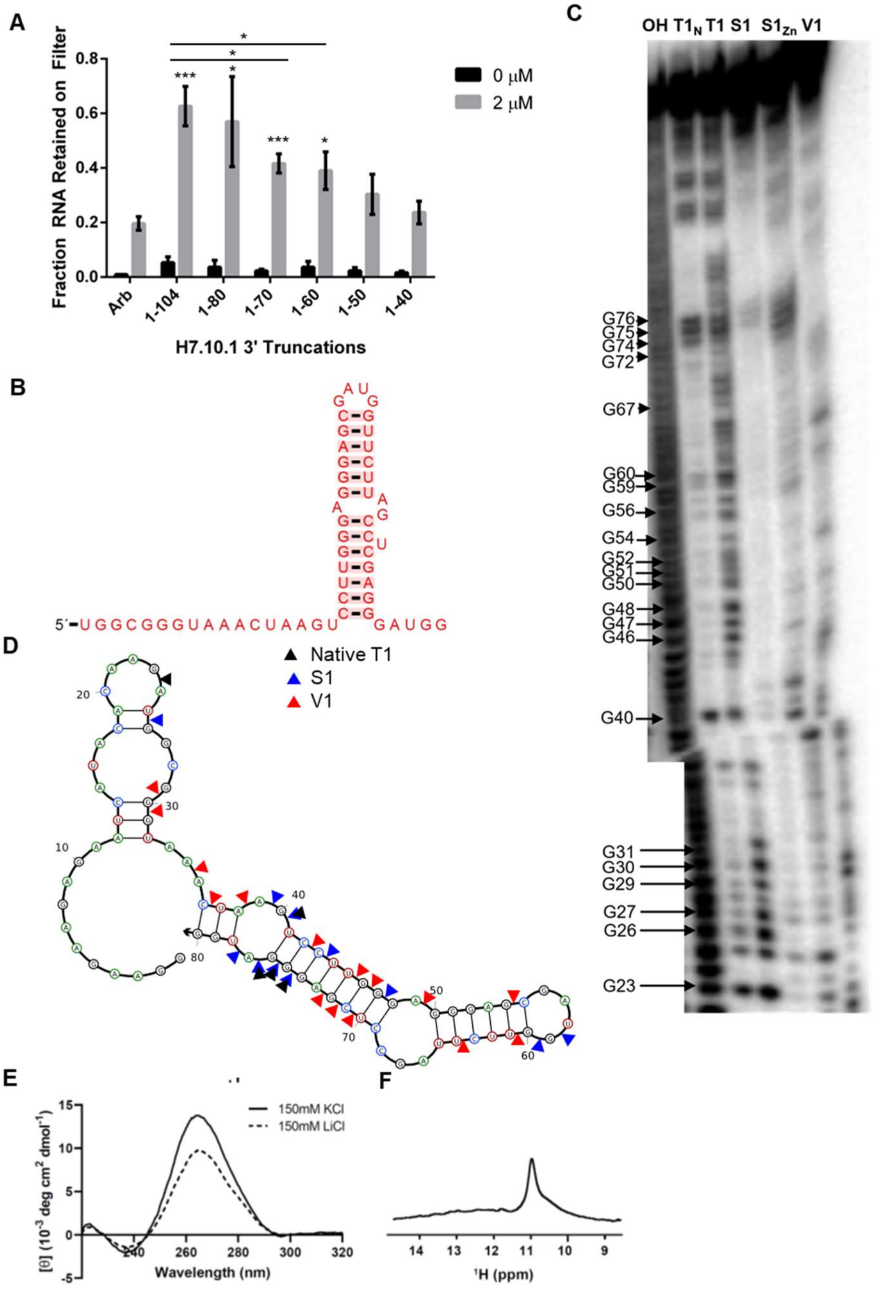


**Figure S22: Exploration of the secondary structure of aptamer H7.10.1.** (A) 3′ truncations of H7.10.1 were assessed for binding to CA lattice using nitrocellulose filter binding assays (n = 3). Values are mean ± SD. * (P value < 0.05), *** (P < 0.001). Statistical comparisons were made versus the level of binding for Arb, except for those against H7.10.1 1-104. (B) Covariance model generated from the sequences from H7.10.1's cluster. The nucleotide identity for all positions was greater than 95% (red letters). Red shading of base pairs refers to no variation within the base pairs (C) Probing of H7.10.1 structure using structure-sensitive nucleases. (D) Mapping the cleavage patterns onto the predicted secondary structure of the 1-80 truncation of H7.10.1. (E) CD spectra of H7.10.1 in 150 mM KCl or LiCl. (F) NMR spectrum of H7.10.1.

**Fig. S23**

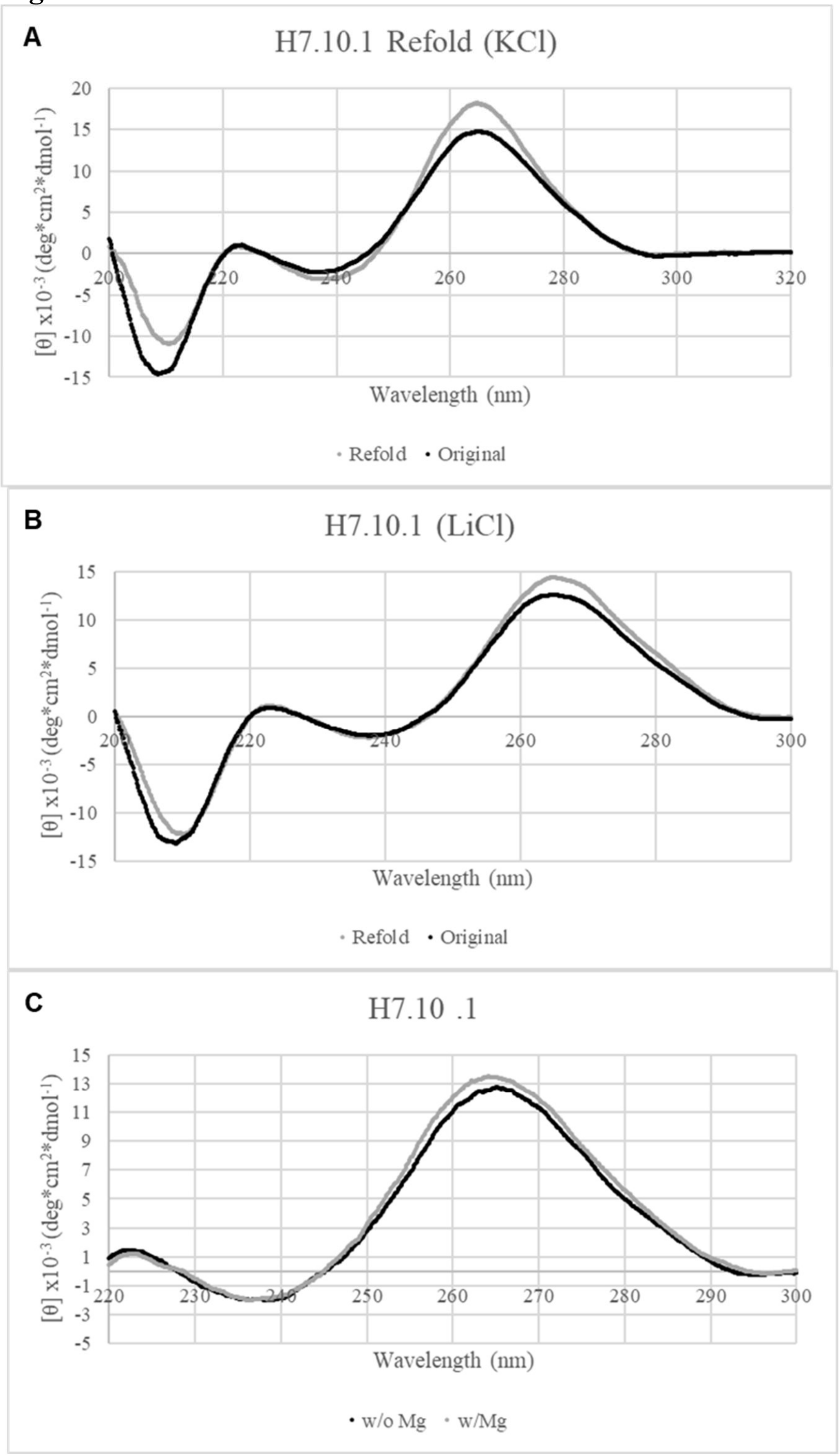


**Figure S23: H7.10.1 Refolding and Magnesium Dependence.** (A) Refolding in KCl. (B) Refolding in LiCl. (C) Magnesium Dependence of H7.10.1.

### Supplemental Tables

**Table S1: Read Counts from Capsid Aptamer HTS Libraries**

| Selection Round | Total Reads | Total Unique Sequences | % Unique Sequences |
|---|---|---|---|
| Lattice Round 8 | 6,118,235 | 2,916,568 | 47.7 |
| Lattice Round 10 | 4,678,445 | 725,600 | 15.5 |
| Lattice Round 15 | 5,449,375 | 150,347 | 2.8 |
| Lattice 15 Nitrocellulose 2 | 3,802,822 | 99,969 | 2.6 |
| Lattice 17 + Hexamer 3 | 4,730,617 | 119,760 | 2.5 |
| Lattice 17 + Hexamer 7 | 3,756,752 | 64,662 | 1.7 |
| Lattice 17 - Hexamer3 | 6,124,384 | 221,093 | 3.6 |

**Table S2: P Values from Statistical Tests of the Aptamer Binding Screen Results**

| Aptamer | P Value 1000 nM Lattice Aptamer vs 0 nM Lattice Aptamer, Significance† | P Value 1000 nM Lattice Aptamer vs 1000 nM Lattice Arb, Significance | P Value 1000 nM Hexamer Aptamer vs 0 nM Hexamer Aptamer, Significance | P Value 1000 nM Hexamer Aptamer vs 1000 nM Hexamer Arb, Significance |
|---|---|---|---|---|
| L15.1.1 | 0.0237, * | 0.0625, ns | 0.4409, ns | 0.7993, ns |
| L15.2.1 | 0.0320, * | 0.0362, * | 0.0177, * | 0.0100, ** |
| L15.3.1 | 0.0190, * | 0.0395, * | 0.2623, ns | 0.1782, ns |
| L15.4.1 | 0.0426, * | 0.0660, ns | 0.1172, ns | 0.3463, ns |
| L15.5.1 | 0.0415, * | 0.1123, ns | 0.1897, ns | 0.4370, ns |
| L15.6.1 | 0.0071, ** | 0.0003, *** | 0.3685, ns | 0.7791, ns |
| L15.7.1 | 0.0696, ns | 0.0904, ns | 0.0742, ns | > 0.9999, ns |
| L15.8.1 | 0.0089, ** | 0.0184, * | 0.5151, ns | 0.5758, ns |
| L15.9.1 | 0.0180, * | 0.0280, * | 0.0576, ns | 0.5232, ns |
| L15.10.1 | 0.0068, ** | 0.0074, ** | 0.3033, ns | 0.9252, ns |
| L15.13.1 | 0.0003, *** | 0.0006, *** | 0.2557, ns | 0.4842, ns |
| L15.14.1 | 0.0446, * | 0.0777, ns | 0.0169, * | 0.2834, ns |
| L15.15.1 | 0.0759, ns | 0.1217, ns | 0.1543, ns | 0.3494, ns |
| L15.16.1 | 0.0005, *** | < 0.0001, **** | 0.2739, ns | 0.6351, ns |
| L15.17.1 | 0.0122, * | 0.0410, * | 0.4204, ns | 0.9227, ns |
| L15.19.1 | 0.0133, * | 0.0253, * | 0.3974, ns | > 0.9999, ns |
| L15.20.1 | 0.0057, ** | 0.0123, * | 0.0600, ns | 0.2566, ns |
| L15.21.1 | 0.0390, * | 0.0777, ns | 0.4382, ns | 0.7972, ns |
| L15.23.1 | 0.0019, ** | 0.0002, *** | 0.5038, ns | 0.9317, ns |
| L15.45.1 | 0.0158, * | 0.0343, * | 0.8714, ns | 0.4235, ns |
| H7.10.1 | 0.0002, *** | < 0.0001, **** | 0.0097, ** | 0.0011, ** |
| H7.12.1 | 0.0059, ** | 0.0136, * | 0.3051, ns | 0.4842, ns |
| H7.20.1 | 0.0065, ** | 0.0136, * | 0.4419, ns | 0.2508, ns |

†ns (P > 0.05), * (P < 0.05), ** (P < 0.01), *** (P < 0.001), **** (P < 0.0001)

**Table S3: Summary of Collected Values**

| RNA | Metal Ion | $Tm_{\alpha = 0.5}$ (°C)[1] | $\alpha_{37°C}$[2] | $\Delta G_{37°C}$ (kcal/mol) | Denatured/Refold | $Mg^{2+}$ Dependence | $^1H$ Imino | Indications of G-quadruplex/tetrad non-canonical base pairing |
|---|---|---|---|---|---|---|---|---|
| L15.6.1 | $Li^+$ | 55.0 | 0.77 | -0.75 | Yes | No | No | No |
| | $K^+$ | 54.8 | 0.78 | -0.79 | Yes | | | |
| L15.7.1 | $Li^+$ | 59.9 | 0.77 | -0.74 | Yes | No | Yes | Yes* |
| | $K^+$ | 61.1 | 0.84 | -1.03 | Yes | | | |
| H7.10.1 | $Li^+$ | 56.4 | 0.80 | -0.86 | Yes | No | Yes | Yes* |
| | $K^+$ | 68.3 | 0.91 | -1.42 | Yes* | | | |
| L15.20.1 | $Li^+$ | 53.0 | 0.76 | -0.70 | Yes | Yes | Yes | Yes |
| | $K^+$ | 58.5 | 0.86 | -1.13 | Yes | | | |

[1]Tm is equal to α at 0.5
[2]α fraction folded at any temperature
* indicates uncertainty

**Table S4: RNA and primer sequences used in this study[a]**

| Name | Sequence |
|---|---|
| L15.1.1 | ggaagaagagaaucauacacaagaGCCGAUGCAGUCCCAAUGUCGUAUGAGAUGUGACUACCUACCACUGUUUAGUGUAUgggcauaagguagguaaguccaua |
| L15.2.1 | ggaagaagagaaucauacacaagaCCACCCAUGCCAAGGAAGGAGGUCGGAGGAGGUCACAGGCAUUGGUGUGCUAGGAUgggcauaagguagguaaguccaua |
| L15.3.1 | ggaagaagagaaucauacacaagaCGCGCAAGCUUUCACGAGGGAGGUUGCGAGGGCGGAUGUGGUUUGCGACGUCGUAgggcauaagguagguaaguccaua |
| L15.4.1 | ggaagaagagaaucauacacaagaCGUGCUCAUACCGCCGAAUACGAGGGAGGAGAGGGUGGCAAAACUGCGUGGGUUgggcauaagguagguaaguccaua |
| L15.5.1 | ggaagaagagaaucauacacaagaCCUACCUGUAUGAAAGAGACGAACCUAACUACAAUGCUAGCAAUGGUGUAUGGUGCgggcauaagguagguaaguccaua |
| L15.6.1 | ggaagaagagaaucauacacaagaUCGACGUACCUCAGGGUGGUGUAUGACUGAGGUGAAGACUGUGAACCAUGGCAUGCgggcauaagguagguaaguccaua |
| L15.7.1 | ggaagaagagaaucauacacaagaCCAACCAGAGACGCACCAGAGUCCUCGAGGGAAAGGAGUGGGUGGUACGGGACAUGgggcauaagguagguaaguccaua |
| L15.8.1 | ggaagaagagaaucauacacaagaCCAUCCCCCAACUUGUCAAGAGGGAAGGAGUGGGAAAGGAAGACAAGGACAUGUGgggcauaagguagguaaguccaua |
| L15.9.1 | ggaagaagagaaucauacacaagaCCCGGCCCCGCCAGUGGAAGUGGGAGGCAUGAGGGAAAGGUUAAACACUGUCGGCUgggcauaagguagguaaguccaua |
| L15.10.1 | ggaagaagagaaucauacacaagaGGACAGAACAACCCUUGAACGAAUGAGGGACAGGUGCGAGGGAAGGAUUCGUUGGAgggcauaagguagguaaguccaua |
| L15.13.1 | ggaagaagagaaucauacacaagaUACCGCGACCUACCCCAGUGCAGGCUAGUGAGGGAGGAGGGUGGCAUCUAGACACUgggcauaagguagguaaguccaua |
| L15.14.1 | ggaagaagagaaucauacacaagaAUCGGAACAAAACGCGACCCGGGACUAGACCUUACCUAAGUGGAUGGGUGUGUGGAgggcauaagguagguaaguccaua |

| | |
|---|---|
| L15.15.1 | ggaagaagagaaucauacacaagaCGCCAAAUUGCUCAGUCCAGUGCGGGAGGUGAGGG CGGAACAACAUGACGGACCAgggcauaagguagguaaguccaua |
| L15.16.1 | ggaagaagagaaucauacacaagaGGCGAAUUACACGACAAUCUGGACGACCUACACUA UGAGAGGAGGGUAGAGUGGAAgggcauaagguagguaaguccaua |
| L15.17.1 | ggaagaagagaaucauacacaagaUGCGCGCGAUGCCCGUACCAUGCUAUGUGGGUGGC GAGGGAGGACACUGCAUGUUGgggcauaagguagguaaguccaua |
| L15.19.1 | ggaagaagagaaucauacacaagaCCGUGCCCAGGGUCUUGCUGUAUGUAUGGUGCUGA UCUACUGUAGCCAUGUGUAUGgggcauaagguagguaaguccaua |
| L15.20.1 | ggaagaagagaaucauacacaagaGGGUCGCAUCGGUCUCUAUGGAUAAAUCUGCCUUA UUGGAGGAUGGGAAAUGUGGAgggcauaagguagguaaguccaua |
| L15.21.1 | ggaagaagagaaucauacacaagaUGGACGAGGUGAUUGUGAUUAGGCUCUACCAGUAU GUCUCCGUUCGUGUAUGGCGUgggcauaagguagguaaguccaua |
| L15.23.1 | ggaagaagagaaucauacacaagaCCCCCCCCAAAGAUACUGGCGCAUGUGAGGGAGGC GAGGGAGGUACGCGUCCUUUgggcauaagguagguaaguccaua |
| L15.45.1 | ggaagaagagaaucauacacaagaAGAGAAUAGGCAAAUACACCAAUCUACCAUAAUGC UGGCGGAGGGAGAAUGAGGAGgggcauaagguagguaaguccaua |
| H7.10.1 | ggaagaagagaaucauacacaagaUGGCGGGUAAACUAAGUCCUUGGGAGGGAGCGAUG GUUCUUAGCCUCGAGGGAUGGgggcauaagguagguaaguccaua |
| H7.12.1 | ggaagaagagaaucauacacaagaGGGUCGCAUCGGUCUCUAUGGAUAACCCUGCCCGG AAGUGGAGGAAUGUGGAgggcauaagguagguaaguccaua |
| H7.20.1 | ggaagaagagaaucauacacaagaUGGACGAGGUGUAUUUGGGCAGACCUUACCUAAGU GGAUGGGUGUGUGGAgggcauaagguagguaaguccaua |
| Arbitrary RNA (Arb) | gggaaaagcgaaucauacacaagaAAUUUGGACUUUCCGCCCUUCUUGGCCUUUAUGAG GAUCUCUCUGAUUUUUCUUGCGUCGAGUUUUCCGGgggcauaagguauuuaauuccaua |
| 15-2 Scrambled | ggaagaagagaaucauacacaagaGGAGGUGCAGGUGCAUAAUGCGUGAAGGCUCGGCCUGUACA CUCGAAGUACUUAGUgggcauaagguagguaaguccaua |
| 56N Forward Primer | GCCTAATACGACTCACTATAGGAAGAAGAGAATCATACACAAGA |
| 56N Reverse Primer | TATGGACTTACCTACCTTATGCCC |
| Arb Forward Primer | GCCTAATACGACTCACTATAGGGAAAAGCGAATCATACACAAGA |
| Arb Reverse Primer | TATGGAATTAAATACCTTATGCCC |
| 56N Reverse + 3′ Tail Extension Primer | TCGTCGTCGTCGTCGTCGTCGTATGGACTTACCTACCTTATGCCC |
| DNA Anti-Tail | TCGTCGTCGTCGTCGTCGTCG |

| | |
|---|---|
| Library-PCR1-F | ACACTCTTTCCCTACACGACGCTCTTCCGATCTGGAAGAAGAGAATCATACACAAG |
| Library-PCR1-R | GACTGGAGTTCAGACGTGTGCTCTTCCGATCTTATGGACTTACCTACCTTATGCCC |
| UDI-PCR2-F | AATGATACGGCGACCACCGAGATCTACAC [i5] ACACTCTTTCCCTACACGACGC |
| UDI-PCR2-R | CAAGCAGAAGACGGCATACGAGAT [i7] GTGACTGGAGTTCAGACGTGTGCTCTTC |

[a]For the RNA sequences, the constant regions are shown in lowercase.

**Table S5: Primer Sequences to Append Illumina Adapters and Sequencing Indices**

| Library | PCR 1 Forward | PCR 1 Reverse | PCR 2 Forward | PCR 2 Reverse | PCR 3 Forward | PCR 3 Reverse[a] |
|---|---|---|---|---|---|---|
| Lattice Round 8 | TAATACGACTCACTATAGGACGCAGGGCTTCATGCACAAGA | TATGGACTTACCTACCTTATGCCC | TTTCCCTACACGACGCTCTTCCGATCTCGCAGGGCTTCATGCACAAGA | GTGACTGGAGTTCAGACGTGTGCTCTTCCGATCTTATGGACTTACCTACCTTA | AATGATACGGCGACCACCGAGATCTACACTCTTTCCCTACACGACGCTCTT | CAAGCAGAAGACGGCATACGAGAT**CGAGTAAT**GTGACTGGAGTTCAGACGTG |
| Lattice Round 10 | TAATACGACTCACTATAGGAAGCATCTATTCATGCACAAGA | TATGGACTTACCTACCTTATGCCC | TTTCCCTACACGACGCTCTTCCGATCTGCATCTATTCATGCACAAGA | GTGACTGGAGTTCAGACGTGTGCTCTTCCGATCTTATGGACTTACCTACCTTA | AATGATACGGCGACCACCGAGATCTACACTCTTTCCCTACACGACGCTCTT | CAAGCAGAAGACGGCATACGAGAT**TCTCCGGA**GTGACTGGAGTTCAGACGTG |
| Lattice Round 15 | TAATACGACTCACTATAGGAAGTAGCTACTCATGCACAAGA | TATGGACTTACCTACCTTATGCCC | TTTCCCTACACGACGCTCTTCCGATCTTAGCTACTCATGCACAAGA | GTGACTGGAGTTCAGACGTGTGCTCTTCCGATCTTATGGACTTACCTACCTTA | AATGATACGGCGACCACCGAGATCTACACTCTTTCCCTACACGACGCTCTT | CAAGCAGAAGACGGCATACGAGAT**AATGAGCG**GTGACTGGAGTTCAGACGTG |
| Lattice 15 Nitrocellulose 2 (L15nc2) | TAATACGACTCACTATAGGAAGAATTGACTCATACACAAGA | TATGGACTTACCTACCTTATGCCC | TTTCCCTACACGACGCTCTTCCGATCTATTGACTCATACACAAGA | GTGACTGGAGTTCAGACGTGTGCTCTTCCGATCTTATGGACTTACCTACCTTA | AATGATACGGCGACCACCGAGATCTACACTCTTTCCCTACACGACGCTCTT | CAAGCAGAAGACGGCATACGAGAT**GGAATCTC**GTGACTGGAGTTCAGACGTG |
| Lattice 17 + Hexamer 3 (L17+H3) | TAATACGACTCACTATAGGACGCAGGGCTTCATGCACAAGA | TATGGACTTACCTACCTTATGCCC | TTTCCCTACACGACGCTCTTCCGATCTCGCAGGGCTTCATGCACAAGA | GTGACTGGAGTTCAGACGTGTGCTCTTCCGATCTTATGGACTTACCTACCTTA | AATGATACGGCGACCACCGAGATCTACACTCTTTCCCTACACGACGCTCTT | CAAGCAGAAGACGGCATACGAGAT**TTCTGAAT**GTGACTGGAGTTCAGACGTG |
| Lattice 17 + Hexamer 7 (L17+H7) | TAATACGACTCACTATAGGAAGCATCTATT | TATGGACTTACCTACCTTATGCCC | TTTCCCTACACGACGCTCTTCCGATCTGCATCTATTCATGCACAAGA | GTGACTGGAGTTCAGACGTGTGCTCTTCCGATCTTATGGACTTACCTACCTTA | AATGATACGGCGACCACCGAGATCTACACTCTTTCCCTACACGACGCTCTT | CAAGCAGAAGACGGCATACGAGAT**ACGAATTC**GTGACTGGAGTTCAGACGTG |

| | | | | | | |
|---|---|---|---|---|---|---|
| | CATGCA CAAGA | | | | | |
| Lattice 17 – Hexamer 3 (L17-H3) | TAATAC GACTCA CTATAG GAAGTA GCTACT CATGCA CAAGA | TATGGA CTTACC TACCTT ATGCCC | TTTCCCTAC ACGACGCTC TTCCGATCT TAGCTACTC ATGCACAAG A | GTGACTGGA GTTCAGACG TGTGCTCTT CCGATCTTA TGGACTTAC CTACCTTA | AATGATACGG CGACCACCGA GATCTACACT CTTTCCCTAC ACGACGCTCT T | CAAGCAGAAGA CGGCATACGAG AT**AGCTTCAG**G TGACTGGAGTT CAGACGTG |

[a]The index sequence is bolded in the PCR 3 reverse primer.
